# Multi-Scale Cortical Bone Traits Vary in Two Mouse Models of Genetic Diversity

**DOI:** 10.1101/2023.06.02.543484

**Authors:** Nicole Migotsky, Surabhi Kumar, John T. Shuster, Jennifer C. Coulombe, Bhavya Senwar, Adrian A. Gestos, Charles R. Farber, Virginia L. Ferguson, Matthew J. Silva

## Abstract

Understanding the genetic basis of cortical bone traits can allow for the discovery of novel genes or biological pathways regulating bone health. Mice are the most widely used mammalian model for skeletal biology and allow for the quantification of traits that can’t easily be evaluated in humans, such as osteocyte lacunar morphology. The goal of our study was to investigate the effect of genetic diversity on multi-scale cortical bone traits of three long bones in skeletally-mature mice. We measured bone morphology, mechanical properties, material properties, lacunar morphology, and mineral composition of mouse bones from two populations of genetic diversity. Additionally, we compared how intra-bone relationships varied in the two populations. Our first population of genetic diversity included 72 females and 72 males from the eight Inbred Founder strains used to create the Diversity Outbred (DO) population. These eight strains together span almost 90% of the genetic diversity found in mice (*Mus musculus*). Our second population of genetic diversity included 25 genetically unique, outbred females and 25 males from the DO population. We show that multi-scale cortical bone traits vary significantly with genetic background; heritability values range from 21% to 99% indicating genetic control of bone traits across length scales. We show for the first time that lacunar shape and number are highly heritable. Comparing the two populations of genetic diversity, we show each DO mouse does not resemble a single Inbred Founder but instead the outbred mice display hybrid phenotypes with the elimination of extreme values. Additionally, intra-bone relationships (e.g., ultimate force vs. cortical area) were mainly conserved in our two populations. Overall, this work supports future use of these genetically diverse populations to discover novel genes contributing to cortical bone traits, especially at the lacunar length scale.

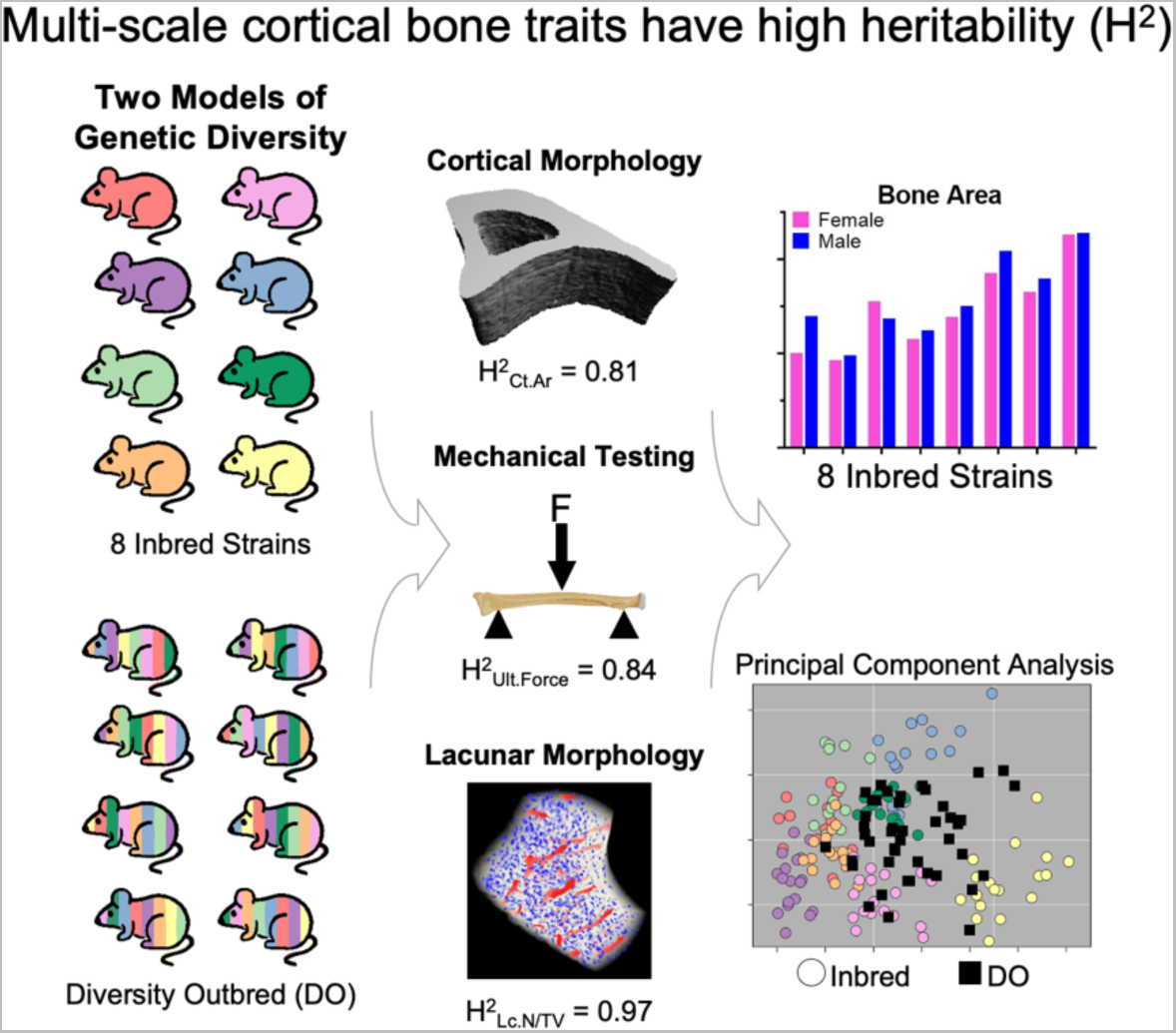

## Introduction

Osteoporosis is a large and growing public health burden that affects more than 54 million people in the United States over the age of 50 and contributes to increased fracture risk ^1–3^. Many individuals are predisposed to osteoporosis or low BMD due to their genetic background. Studies of twins and generations of sisters have shown that about 70% of variability in bone density and about 60% of variability in osteoporotic fracture risk is genetically based ^4–6^. Clinically, osteoporosis is defined using bone mineral density (BMD) from DEXA scans, however many traits in addition to BMD contribute to bone strength and fracture risk, such as morphology, mineralization, and stiffness^7^; it is important to understand if these bone traits are also heritable.

Classically, heritability is estimated using twin, adoption, or family clinical study designs^8, 9^. A review of heritability studies of body mass index using twins from 12 different countries showed high variability between studies and that heritability is sensitive to age, GDP, and economic growth rates showing the importance of considering environmental factors in addition to genetic factors^10^. Karasik et al. reported heritability of bone morphology as high as 98.3% (tibia cortical area fraction) estimated from the Framingham Offspring study participants^11^. With the increase in availability and decrease in cost of whole-genome arrays, heritability estimates from genome-wide association studies (GWAS) have become more standard^8^. Due to the high homology between mammalian genomes, results from mouse studies can assist in understanding the complicated role of genetics in humans^9^. Additionally, environmental factors can be controlled in mice and more complex phenotypes can be measured (e.g., bone strength or lacunar morphology). A recent study in mice by Al-Barghouthi et al. showed that all 55 skeletal phenotypes measured in 12-week-old genetically diverse mice had non-zero heritability^12^. Understanding the genetic basis of cortical bone traits in adult skeletons can allow for the discovery of novel genes or pathways as therapeutic targets for low bone mass^13^.

There remain gaps in the current literature regarding the heritability of bone traits. First, almost 80% of all GWAS participants are of European decent despite only accounting for approximately 16% of the world population ^14^. Likewise, the bone data collected in the Framingham Study is only from people European decent^11^, limiting the applicability of results to broader populations. Second, previous mouse studies only investigated a single long bone, primarily the femur^12, 15, 16^. Many bone traits are skeletal site dependent, so heritability of traits may also vary between long bones. Third, measured traits also focused on the whole-bone or tissue length scale, but none have investigated the tissue or cellular length scale. Finally, phenotyping of and heritability measurements for cancellous or cortical bone was previously done using skeletally-developing mice (11-13 week old) or elderly humans (avg 72 years old), leaving a gap to investigate the heritably of traits in the young-adult, skeletally-mature skeleton, i.e. when bone mass is maximized. Bone mass and fracture risk later in life depends on the peak bone mass attained as a young-adult; a 10% increase in peak bone mass reduces fracture risk in older adults by 50%^17^ making it important to study what factors affect the young-adult skeleton.

To facilitate the investigations of the genetic basis of various diseases and phenotypes, The Jackson Laboratory (JAX) generated the Diversity Outbred (DO) mouse population by cross-breeding eight inbred founder strains to produce a population with random assortments of genetics modeling the heterozygosity of the human population and the wide range of human diseases^18, 19^. These eight inbred strains consist of three wild-derived strains (CAST/EiJ, PWK/PhJ, and WSB/EiJ) and five classical laboratory strains (A/J, C57BL/6J, 129S1/SvImJ, NOD/ShiLtJ, and NZO/HlLtJ) that together cover almost 90% of the genetic diversity found in the mouse genome and represent the three sub-species of the common house mouse (*Mus musculus domesticus, Mus musculus musculus,* and *Mus musculus castaneous*)^20^. Over the last decade, use of these genetically diverse populations has increased in the bone field, and these mice have been used to evaluate the heritability and candidate genes regulating cancellous bone microarchitecture in the growing skeleton^16^, the response to hindlimb disuse via casting of one femur^15^, and the heritability of femur properties and genes influencing cortical bone accrual in the growing skeleton^12^. These studies provide motivation and rationale to use these populations of genetically diverse mice, both the Inbred Founders and DO, to study the heritability of cortical bone traits.

In this study, we aimed to calculate the heritability of multi-scale cortical bone traits of the three long bones – the radius, tibia, and femur – in skeletally-mature mice. We used two models of genetic diversity with each population containing the same pool of possible alleles. First, a cohort of males and females from the eight Inbred Founder strains used to create the Diversity Outbred (DO) population, and second, a cohort of males and females from the DO population. Comparing the individual inbred strains, we hypothesized that cortical bone traits vary with genetic background and sex. Comparing the Inbred Founder cohort to the DO cohort, we hypothesized the DO cohort has a similar spread and mean per trait as the Inbred Founder cohort. Comparing the correlations between traits within a single animal, we hypothesized that relationships between cortical bone traits are conserved in these two models.

## Methods

### Mouse Populations

All mouse work was completed with approval of the Washington University Institutional Care and Use Committee (IACUC). Two mouse populations were used to model genetic diversity: (1) Eight Inbred Founder strains and (2) Diversity Outbred (DO) mice from JAX. The eight Inbred Founder strains (CAST/EiJ – JAX stock #000928, PWK/PhJ - #003715, WSB/EiJ - #001145, A/J - #000646, C57BL/6J - #000664, 129S1/SvImJ - #002448, NOD/ShiLtJ - #001976, and NZO/HlLtJ - #002105) have been continuously outbred by Jackson Laboratory to create and maintain the Diversity Outbred population (#009376) (Fig 1A). Inbred Founder strains (n = 9/strain/sex) were delivered at 8 wks of age and aged to 22 wks in our animal facilities. Of the N = 144 that arrived, eight were lost (3 NOD and 5 CAST) leaving N = 136 for analysis. DO mice (n = 25/sex, G46 and G47) were delivered between 3 and 5 wks of age and aged to 22 wks in our animal facilities. One female DO mouse died prematurely leaving 24 for analysis. Inbred mice were group housed up to 5 in a cage. Male DO mice were housed individually, and female DO mice were housed up to 3 in a cage according to recommendations by JAX to reduce in-fighting. All mice were kept on a 12 hr light-dark schedule with *ad libitum* access to food and water.

**Figure 1:**
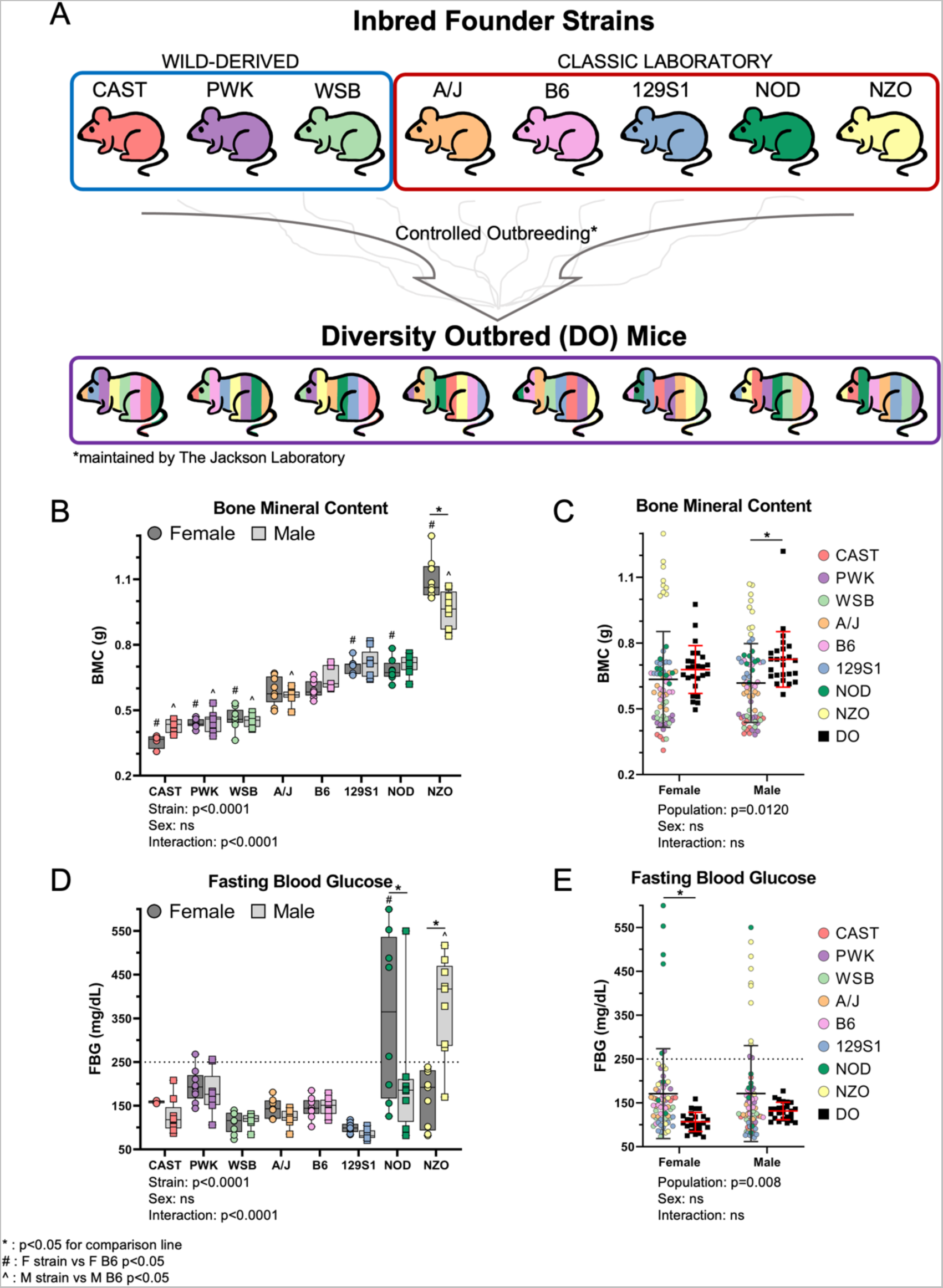
A) Two mouse models of genetic diversity were used, eight inbred mouse strains and an outbred stock. The inbred strains were the founders used by The Jackson Laboratory to create the diversity outbred (DO) population. After controlled outbreeding of the founder lines, each DO mouse is genetically unique. Phenotyping was performed on 5 mo-old females and males of the eight Inbred Founder strains and DO mice from generations 46 and 47. B) Whole-body bone mineral content of the Inbred Founders shows a 3-fold difference in BMC between the largest (NZO F) and smallest (CAST F) mice. Strains on the x-axis are ordered from smallest to largest BMC. This order is maintained for all graphs in the study. C) The DO mice have a larger average BMC with less variation than the pooled Inbred Founder population. D) Fasting blood glucose shows that about 12% of the Inbred Founder mice become hyperglycemic (FBG > 250mg/dL), mostly NOD F and NZO M. E) All DO mice have FBG levels in the healthy range. *p<0.05 for comparison line; # p<0.05, F strain vs F B6; ^p<0.05, M strain vs M B6

### Longitudinal Measurements

After one week of acclimation, Inbred Founder strains were monitored for whole body growth. Mouse weights, dual-energy x-ray absorptiometry (DEXA) scans, and fasting blood glucose (FBG) were collected at 9, 12, 15, 18, and 22 wks of age. Briefly, mice were fasted for 6 hrs, weighed, and FBG was measured using a drop of blood from the tail (Glucocard Vital, Arkray Inc.) to track glycemic status; hyperglycemia was defined as a level over 250 mg/dL. Mice were then anesthetized with isoflurane (1.5-4%) for 10-15 minutes to ensure full sedation during the DEXA scan. Whole-body (excluding the head) bone mineral content (BMC) and areal bone mineral density (aBMD) were measured from DEXA (UltraFocus, Faxitron, 4 x 40 kV and 6 x 80 kV scans). At 22 wks old, Inbred Founder mice were euthanized, bones collected for analysis (Table 1), and bodies stored frozen at -20°C.

**Table 1:** Bones evaluated for various outcomes in both the Inbred Founders and Diversity Outbred (DO) mice.

| Bone | Outcome |
| --- | --- |
| Left Tibia | XRM |
| Right Tibia | μCT (Supp) |
| Left Femur | μCT (Supp) |
| Right Femur | Raman |
| Right Radius | μCT - 3pt bend |

Diversity Outbred Bone Outcomes
| Bone | Outcome |
| --- | --- |
| Right Tibia | μCT (Supp) |
| Left Femur | μCT (Supp) |
| Right Radius | μCT - 3pt bend |

DO mice were evaluated *in vivo* at 22 wks. Briefly, mice were fasted for 6 hrs, weighed, and FBG was measured. The next day mice were anesthetized with isoflurane for DEXA and subsequent tibial microCT (µCT) (*see Cortical Bone Morphology*). DO mice were subjected to right limb tibial loading (findings not reported here), euthanized at 25 wks age, femora and radii collected for analysis (Table 1), and bodies stored frozen at -20°C. Only tibial µCT from before loading in the DO mice were analyzed in this study. No Inbred Founder mice were loaded.

### Cortical Bone Morphology

The right tibia was µCT scanned (vivaCT 40, Scanco Medical, Switzerland - 70kVP, 8W, 300ms integration time) at 10.5 µm/pixel resolution either *ex vivo* (Inbred Founders) or *in vivo* (DO mice). A 1.05 mm long region (100 slices) centered 5 mm proximal to the distal tibiofibular junction (TFJ) was analyzed.

The entire right radius was µCT scanned (µCT50, Scanco Medical, Switzerland-70kVP, 4W, 700ms integration time) at 7.4 µm/pixel resolution *ex vivo*. A 1.48 mm long region (200 slices) centered at the radius midpoint was analyzed.

The left femur was µCT scanned (µCT50, 70kVP, 4W, 700ms integration time) at 7.4 µm/pixel resolution *ex vivo*. A 1.11 mm long region (150 slices) centered at the femur midpoint was analyzed.

All bones were analyzed following published guidelines^21^ for total area (Tt.Ar), bone area (Ct.Ar), medullary area (Ma.Ar), cortical thickness (Ct.Th), polar moment of inertia (pMOI), and tissue mineral density (TMD).

### Mechanical Testing

The right radius was cleaned of all tissue and stored at -20°C in PBS-soaked gauze until testing. The radius was selected for mechanical testing because its relatively long, slender shape allows for more accurate estimation of material properties than the femur^22^. On the day of testing, samples were brought to room temperature and kept hydrated in PBS. Bones were tested in three-point bending with a bottom span of 7 mm and the top point aligned with the bone center and held in place with a -0.1 N pre-load. Bones were pre-conditioned for 5 cycles at an amplitude of -0.4 N at 1Hz then loaded monotonically to failure at 0.1 mm/s. Force-displacement curves were analyzed for stiffness (K), ultimate force (F_u_), yield force (F_y_), post-yield displacement (PYD), and work-to-fracture (Wfx). Material properties (ultimate stress (S_u_), yield stress (S_y_), Young’s elastic modulus (E)) were estimated using engineering beam theory^23^.

### Osteocyte Lacunar Morphology

Osteocyte lacunar morphology was quantified in the Inbred Founder population only. The left tibiae of Inbred Founders were dissected and fixed in 4% PFA overnight, then rinsed and stored in PBS until use. Bones were cut transversely 5 mm proximal to the TFJ using an Isomet saw, and the distal portion was scanned on an X-ray microscope (XRM, Xradia Versa 520, Zeiss) 4 mm proximal to the TFJ. A pre-scan, using the 4x objective, was acquired to locate the high-resolution scan region of interest, centered halfway between endocortical and periosteal surfaces at the postero-lateral apex (Supp. Fig. S8A). Scanning parameters for the high-resolution scan were: 40 kV, 3 W, 1601 projections, 20x objective, bin = 2, 4800-5000 projection intensity (7-8 sec exposure), yielding a nominal resolution of 0.54 µm/voxel. Bones were segmented, filtered, and analyzed using custom scripts in Dragonfly (version 4.1, Object Research Systems, Montreal, Canada) for total lacunar volume, total vessel volume, and individual lacunar properties (volume, aspect ratio, phi, and sphericity), as described^24^.

### Raman Spectroscopy

Bone matrix composition was quantified in cortical bone from the Inbred Founder population only. The right femurs of Inbred Founders were dehydrated in ethanol and embedded in poly methyl methacrylate (PMMA) (Thermo Scientific AAA130300F, Thermo Fisher Scientific). Plastic blocks were cut at the midpoint perpendicular to the bone long-axis and trimmed to 5 mm in length. Blocks were polished using an Allied TechPrep polisher to 0.05 µm (600 grit silicon carbide sandpaper, 1200 grit silicon carbide sandpaper, 1 µm aluminum oxide, 0.05 µm aluminum oxide) on Rayon felt pads (Allied). Measurements were taken on the anterior side of the femur in lamellar bone excluding the periosteal and endocortical surfaces (Supp. Fig S1). Measurements were taken in a grid of 3 spots wide spanning the cortical width (5-9 spots long) spaced 20 µm apart. Data was collected using a red laser (785 nm wavelength) on a Renishaw inVia confocal microscopy system (Renishaw, Wotton-under-Edge, Gloucestershire, UK). Each point was collected with 10 accumulations with a 6 second exposure time and post-processed using WiRE 4.1 software (baseline subtraction with 11^th^ order polynomial fit, cosmic ray removal, spectra normalization). A single measurement of the PMMA for each sample was used for baseline subtraction. All data was analyzed in a custom MATLAB code to quantify area ratios of mineral:matrix (v2 phosphate: amide III, v1 phosphate:proline), carbonate:phosphate (v1 carbonate: v1 phosphate), and crystallinity (inverse of full-width at half-maximum of v1 phosphate)^25–27^.

### Correlation Matrix

A matrix of Pearson’s correlation coefficients was computed using R (v4.0.2) on three separate datasets: 1) all lacunar traits in the Inbred Founders, 2) radial and whole body traits in the Inbred Founders, and 3) radial and whole body traits in the DO mice. Correlations were calculated using the *cor* function and visualized using the *corrplot* function. For the lacunar traits, variables were unbiasedly hierarchically clustered into three groups using the *ward.D2* algorithm. For the radial traits, variables were ordered by measurement technique consistently for each mouse population (Inbred Founders and DO).

### Principal Component Analysis (PCA)

Principal component analysis (PCA) was performed using R on two datasets from the Inbred Founder population: 1) radial and whole-body traits (15 traits), and 2) lacunar traits (10 traits). Each trait was centered and scaled within the *prcomp* function to have a mean of 0 and standard deviation of 1. The contribution of each trait to each principal component (PC) was extracted from the PCA. To compare the Inbred Founders and DO population for dataset 1 (radial and whole-body traits), each DO animal was plotted onto the PCA space defined by the Inbred Founders. Briefly, the value for each DO trait was centered and scaled and linearly combined according to PC weightings per variable from the Inbred Founder PCA. For visualization of the Inbred Founders, data points were grouped by mouse strain and encompassed by a normal data ellipse spanning one standard deviation (68%).

### Heritability Calculations

Broad-sense heritability (H^2^) was calculated for all traits measured in the Inbred Founder mice as the proportion of variance due to genetic differences according to Moran et al.^28^. Heritably was calculated as H^2^ = σ^2^_strain_/(σ^2^_strain_+ σ^2^_sex_+ σ^2^_res_) where σ^2^_strain_ is the between-strain variance, σ^2^_sex_ is the between-sex variance, and σ^2^_res_ is the residual variance (equation on Table 2). Variances were calculated from the sum of squares from a 2-factor analysis of variance (ANOVA) with strain and sex and the two factors. Specifically, σ^2^_strain_ = SSstrain/navg (navg = average group size), σ^2^_sex_ = SS_sex_/df_sex_, and σ^2^_res_ = SSres/df_res_.

**Table 2:**
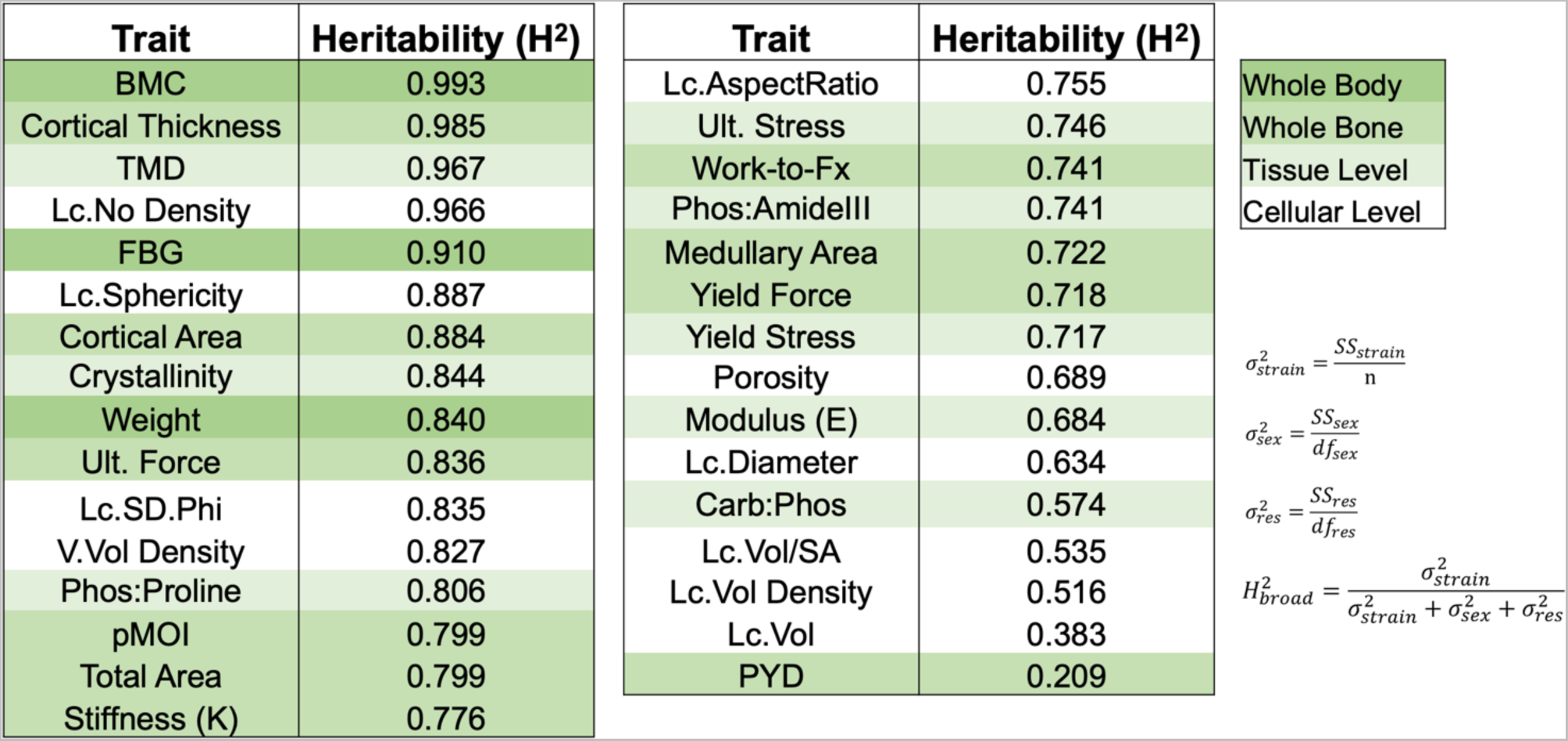
Broad sense heritability was calculated using data for the Inbred Founders for each trait as the fraction of total variation due to strain using the equation shown.

### Statistical analysis

Statistical analysis was done in GraphPad Prism (v9). First, outcomes from the eight Inbred Founders were analyzed using a 2-factor ANOVA with mouse strain and sex as the two factors. Second, outcome from the Inbred Founder population (all eight strains pooled) and the DO population were compared using a 2-factor ANOVA with population and sex as the two factors. Post-hoc tests were run with Sidak correction. Significance was set at p < 0.05. Body mass was evaluated as a co-variate using a full factorial general linear model in SPSS (IBM, Armonk, NY). Body mass adjusted values per animals were calculated according to guidelines in Jepsen et al^23^ using the following equation *Trait_adjusted_ = Trait_unadjusted_ − slope \** (*body mass − mean body mass*). Slopes were determined from a linear regression of each trait with body mass within each strain group (8 groups). To ensure statistically meaningful relationships, for any regression with a p > 0.20 the slope was set to 0 for that strain. The mean body mass was calculated as the mean of the means of each strain group (mean of 8 means).

## Results

Young-adult mice from the Inbred Founder and Diversity Outbred populations span a large range of body size and bone mass At the whole-body length scale, DEXA was used to assess skeletal mass. The eight Inbred Founder strains were skeletally mature by 22 wks; the average increase in BMC between 18 and 22 wks is less than 4% of final BMC (0.2% - 7.7%) (Supp Fig S2A). At 22 wks, BMC of the Inbred Founders varied significantly between strains, with a 3-fold difference between CAST females and NZO females (Fig 1B). (Note that Inbred Founder strains are ordered from smallest to largest BMC along the x-axis; this order is maintained for all graphs in this report.) The main effect of sex on BMC was not significant, although there was a significant strain-sex interaction, driven by a greater BMC in NZO females than males (Fig 1B). The DO population had significantly higher BMC than the Inbred Founder population with a narrower range of values and higher minimum values (Fig 1C). This result was more pronounced in the males. Similar trends across strains and between the two populations were seen with body weight (Supp Fig S2 D,E). About 12% of the Inbred Founders were hyperglycemic by 22 wks (Fig 1D). NZO males became hyperglycemic between 12 and 15 wks of age (Supp Fig S2B), which coincided with a halt in an increase in their BMC and weight (Supp Fig S2 A,C). NOD females became hyperglycemic between 18 and 22 wks of age (Supp Fig S2B), which coincided with a loss in their weight (Supp Fig. S2C). In contrast, none of the DO mice were hyperglycemic at 22 wks (Fig 1E).

### Cortical morphology of three different long bones varies with mouse strain and sex

Whole-bone morphology, assessed using µCT, varied significantly between Inbred Founder strains in a sex-dependent manner (Fig 2 and Supp Fig. S3). Radial bone area (Fig 2A) spanned a 1.7-fold difference between smallest (PWK females) and largest (NZO males), while medullary area (Fig 2B) spanned a 3.9-fold difference between smallest (WSB females) and largest (NZO males). Bone area was positively correlated with skeletal mass and body weight (r=0.85 and 0.75 respectively, Fig 4A). While all strains contributed to these overall trends, WSB mice had larger bones and NOD mice had smaller bones than strains of similar BMC (Fig 2A). Compared to the Inbred Founders, the DO population on average had significantly larger bones with a similar medullary area (Fig 2 D,E). Analysis of morphology from tibias (Supp Fig S5) and femurs (Supp Fig S6) showed similar population spreads and differences between mouse strains as observed for the radius, indicating consistency between long bones. Radius tissue mineral density also varied significantly between Inbred Founder strains, although there was no main sex effect or sex-strain interaction (Fig 2C). The DO males, on average, had a lower TMD than Inbred Founder males, but no difference was observed between females (Fig 2F). Similar results were seen in the femoral TMD (Supp Fig S6K). However, in the tibia, the DO population had higher TMD as compared to the Inbred Founders, especially in females (Supp Fig S5K). In summary, cortical morphology varied widely between Inbred Founder strains with outbred animals (DO) having larger bones on average.

**Figure 2:**
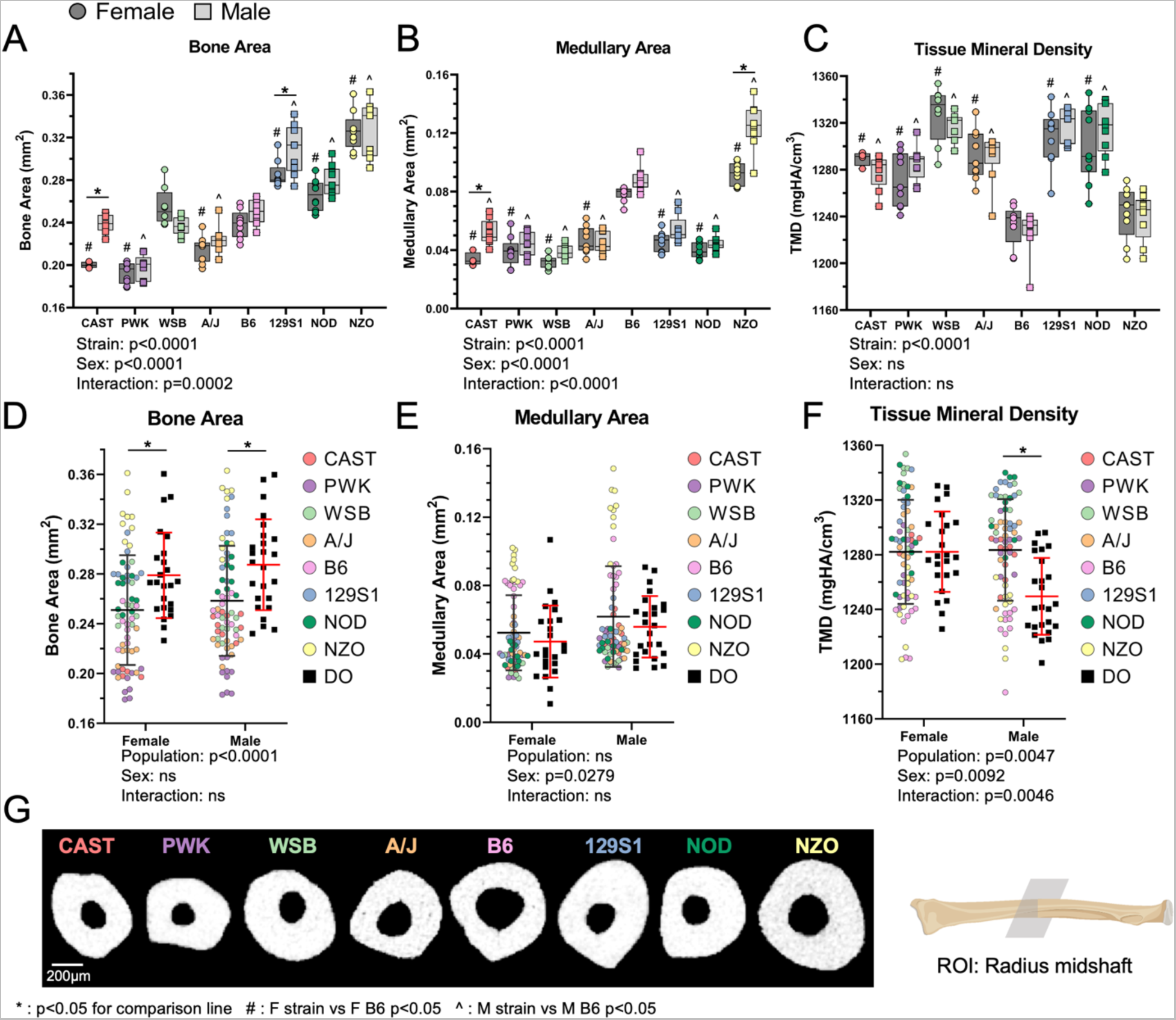
Radial morphology varies with strain in a sex-dependent manner. A) Bone area and B) medullary area of Inbred Founder mice have significant strain, sex, and strain-sex interaction terms (ANOVA). C) Tissue mineral density shows significant strain differences in the Inbred Founder mice. D) The DO population has higher bone area than the pooled Inbred Founder population, but no difference in E) medullary area. F) The male DO population has lower TMD than the male Inbred Founders. This difference is not seen in females. G) Representative cross-sections at the radius mid-diaphysis show the variation in bone shape and size between the eight Inbred Founder strains (females). *p<0.05 for comparison line; # p<0.05, F strain vs F B6; ^p<0.05, M strain vs M B6

Cortical bone mechanical properties vary with mouse strain and sex, but material properties only vary with mouse strain The radius was tested in three-point bending to determine whole-bone mechanical and estimated material properties (Fig 3A). Only the radius was selected for testing since it closely fits the assumptions of beam theory, allowing for accurate estimation of material properties^22^. In the Inbred Founder population, all mechanical properties varied between strains with significant strain, sex, and strain-sex interaction effects (Fig 3 B,D and Supp Fig. S4 A-C). Ultimate force, a measurement of whole-bone strength, spanned a 2.7-fold difference between the weakest strain (PWK females) and the strongest (NZO males) (Fig 3B). Mouse strain accounted for about 80% of the total variation in ultimate force (ANOVA). Compared to the Inbred Founders, the DO population had stronger bones on average, and no DO mice had bones as weak as any of the weakest Inbred Founders (PWK) (Fig 3C). Post-yield displacement (PYD), a measurement of ductility, varied only modestly with strain (Fig 3D) and did not correlate strongly with BMC (r=0.24, Fig 4A). The radius from B6 females were the least ductile and those from B6 males the most. PYD was highly variable within each strain/sex group and mouse strain only accounted for about 11% of the total variation. The PYD in the DO population did not differ from the PYD in the Inbred Founder population (Fig 3E). All estimated material properties varied between Inbred Founder strains but did not differ between sexes. Ultimate stress had a 1.3-fold difference between PWK males (lowest) and WSB females (highest) (Fig 3F). The DO population had slightly lower ultimate stress than the Inbred Founder population (Fig 3G), which is also true of other material properties (Supp Fig S4 I,J). In summary, radial bone mechanical and material properties vary between the Inbred Founder strains, with the outbred mice (DO) having average higher mechanical but lower material properties than the average Inbred Founder.

**Figure 3:**
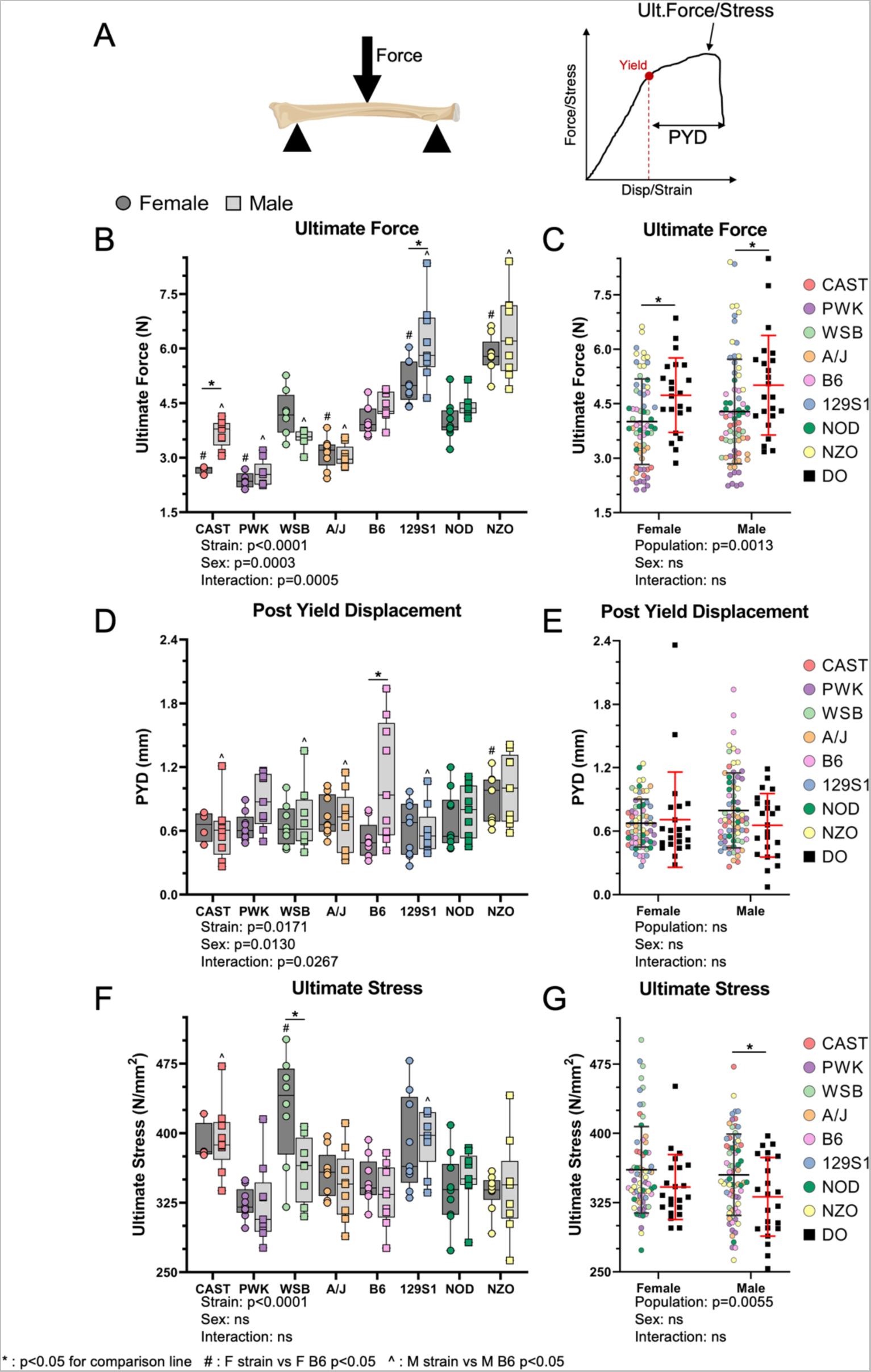
Radius mechanical and material properties vary between inbred mouse strains. A) The radius was tested using three-point bending to determine mechanical properties. Beam theory equations were used to estimate material properties. B) Ultimate force varies between Inbred Founder strains with significant strain, sex, and strain-sex interaction. C) The DO population has stronger bones (higher ultimate force) than the pooled Inbred Founder population. D) Post-yield displacement (a measure of ductility) also had significant strain, sex, and strain-sex interactions. E) The ductility of DO population is not significantly different from the Inbred Founders. F) Ultimate stress and other material properties varied with mouse strain with no significant sex differences. G) The DO population has lower material properties, such as ultimate stress, than the Inbred Founder population. *p<0.05 for comparison line; # p<0.05, F strain vs F B6; ^p<0.05, M strain vs M B6

**Figure 4:**
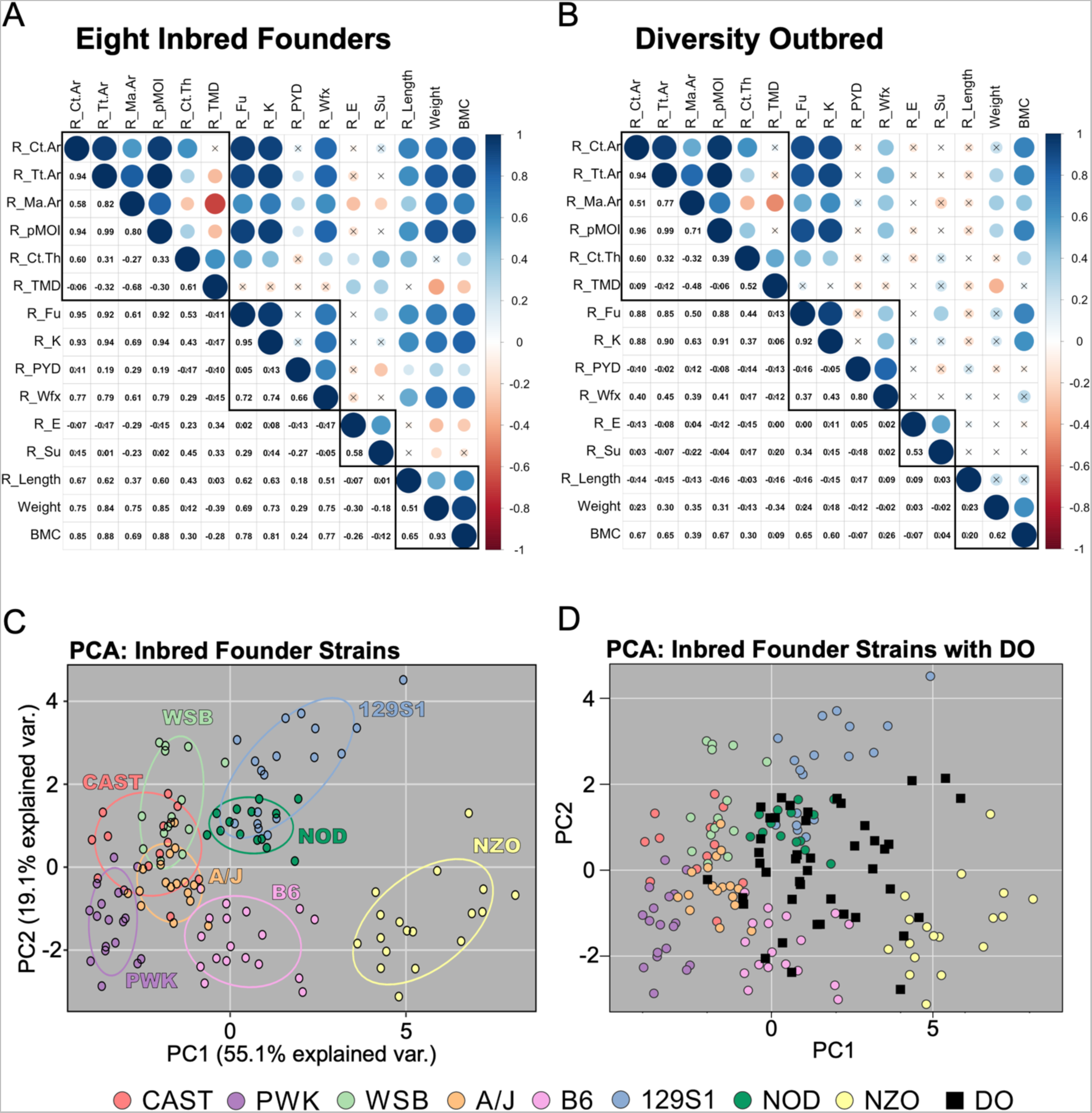
A) Matrix of Pearson’s correlations between traits measured in the radius in the eight Inbred Founder strains (A) and the Diversity Outbred mice (B). Black boxes separate different types of outcomes (morphology, mechanical properties, material properties, whole skeleton size). Similar correlations are seen between the two populations, but are generally weaker in the DO population. For A) and B) an X indicates a non-significant correlation (p > 0.05) C) PCA analysis showing how the Inbred Founders cluster by strain. D) The DO mice mapped onto the PCA space defined by the Inbred Founders; the DO population lays in a subset, hybrid space eliminating extreme values at the periphery of the PCA space. Ct.Ar: Cortical Area, Tt.Ar: Total Area, Ma.Ar: Medullary Area, Ct.Th: Cortical Thickness, TMD: Tissue Mineral Density, Fu: Ultimate Force, K: Stiffness, PYD: Post-yield displacement, Wfx: Work to fracture, E: Young’s Elastic Modulus, Su: Ultimate Stress, BMC: Bone Mineral Content

Within-bone correlations are similar in both mouse populations, but stronger in the Inbred Founder population

Our previous work^29^ showed that cortical bone traits are highly correlated within a single bone (femur or radius) in an Advanced Intercross mouse population (LGXSM). Here, we investigated the relationship between traits by calculating the Pearson’s correlation coefficient for 15 whole-body and radial trait pairs in the Inbred Founder and DO populations (Fig 4). In the Inbred Founders, bone size traits (bone area, total area, medullary area, and pMOI) were all highly correlated with each other (Fig 4A). Cortical thickness was only moderately correlated with other morphology parameters. Bone size traits were highly correlated with whole-bone mechanical properties (with exception of PYD) as well whole-body measurements (bone length, weight, BMC). Mechanical properties (except PYD) were also correlated with whole-body measurements. Bone material properties correlated with each other, but did not correlate with bone size parameters, which is expected because material properties are already normalized for bone size. Tissue mineral density had few strong correlations but was moderately correlated with cortical thickness (r = 0.61) and medullary area (r = -0.68). Bones with high tissue mineral density tend to have a thick cortex and a small medullary cavity.

Compared to the Inbred Founders, the DO population showed similar correlations between traits, but the magnitudes were generally weaker. In particular, correlations with body weight and length were much lower in DO mice (Fig 4B). The correlation coefficient between total area and BMC was lower by 0.23 (0.88 in Founders vs 0.65 in the DO), while the correlation between total area and weight was lower by 0.54 (0.84 in the Founder vs 0.30 in the DO). Additionally, the direction (negative or positive) of many correlations with bone length were opposite in the two populations. The relationship between ultimate force and bone area was highly, positively correlated (r=0.95 in Inbred Founders and 0.88 in DO) and extremely conserved in the two populations, sharing the same slope and intercept (Supp Fig S9).

Inbred Founder strains separate using PCA while Diversity Outbred mice overlap many individual strains and occupy gaps between strains

PCA was done to evaluate within and between strain differences based on multiple (15) traits from the radius of the Inbred Founders (Fig 4C). The first two principal components (PC) explained almost 75% of the population variance, and mice strongly clustered by strain. Mouse and bone size and strength parameters (BMC, weight, Ct.Ar, Tt.Ar, pMOI, Fu, K) were the main contributors to PC1. Bone thickness and material properties (Ct.Th, TMC, Su, E) were the main contributors to PC2 (Supp Fig S7C). NZO was separate from all other strains, while B6 was mostly separate (small overlap with A/J only). 129S1 and NOD showed moderate overlap. The wild-derived strains showed moderate overlap with each other (CAST overlaps WSB and PWK), and with A/J. When the DO mice were mapped onto the PCA space defined by the Inbred Founders, they occupy a smaller, central region of the PCA space, eliminating extreme values at the periphery (Fig 4D). Additionally, many DO mice filled the empty space between the NZO and 129S1 clusters indicating that individual DO mice have a unique combination of bone traits and do not necessarily phenotypically mimic a single Inbred Founder mouse.

### Lacunar morphology varies between Inbred Founder strains

Osteocyte lacunar morphology was analyzed in the tibia of Inbred Founder mice using high-resolution XRM (Fig 5 and Supp Fig S8). Over 4,000 lacunae were individually analyzed per sample. Total lacunar number density (Lc.Num/TV) varied significantly between strains in a sex-dependent manner ranging from 53,189/mm^3^ in B6 females to 80,200/mm^3^ in 129S1 females (Fig 5B). The total volume of lacunae only occupied 1-2% of the total bone volume (cortical bone tissue including pores), but varied significantly between Inbred Founders (Fig 5C). Quantifying approximately 4,000 lacuna per mouse, the median lacunar volume also varied with mouse strain in a sex-dependent manner (Fig 5D). Lacunar porosity (Lc.Vol/TV) highly, positively correlated to median lacunar volume (r = 0.65), and moderately correlated with lacunar number density (r = 0.49), indicating that the total lacunar volume is increased mainly by increasing the volume of each lacuna rather than the number of lacunae (Fig 5G).

**Figure 5:**
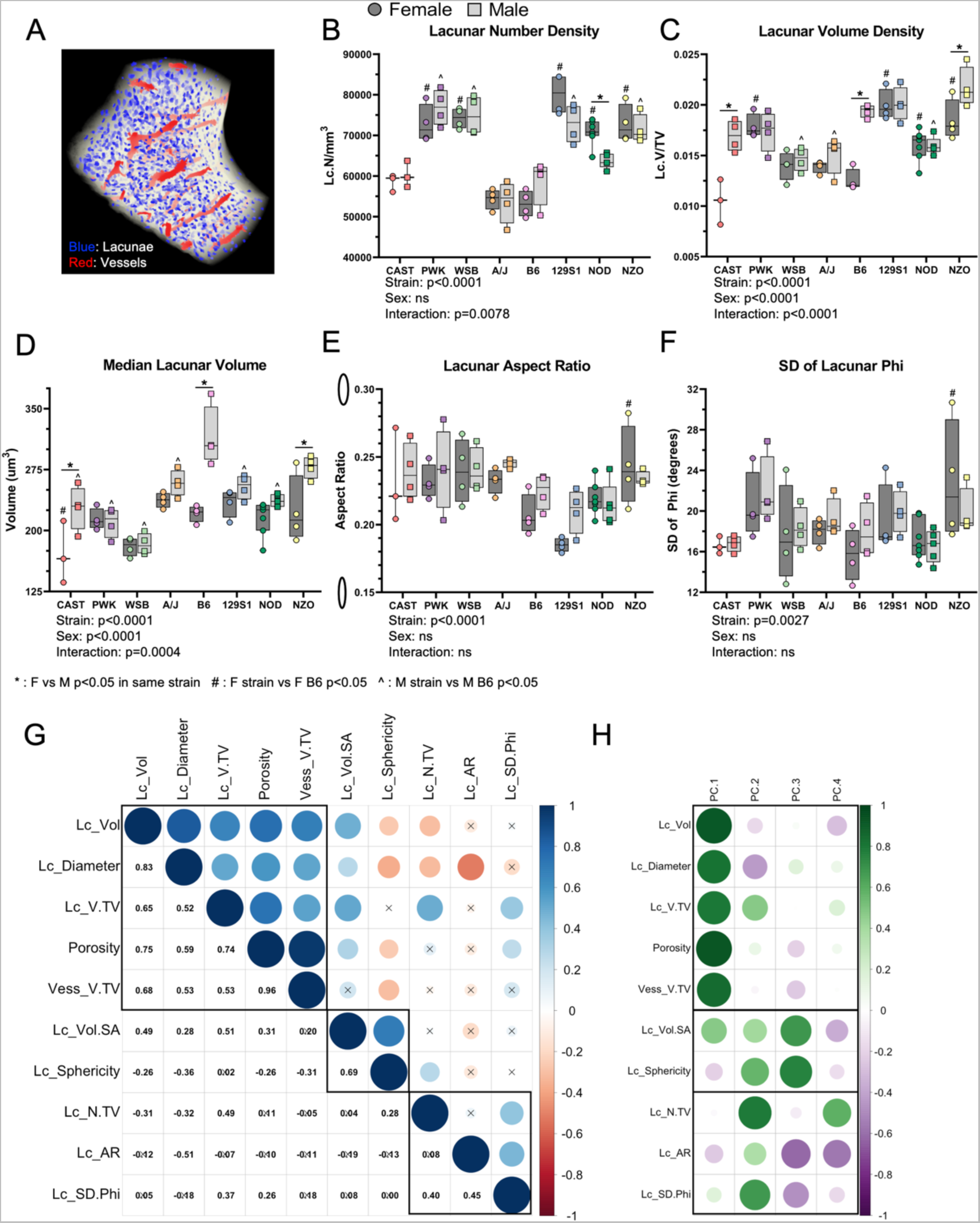
Osteocyte lacunar morphology varies between inbred strains. A) Lacunae were imaged using x-ray microscopy (XRM). Representative cross-section of the field of view (FOV) imaged. Intra-cortical pores were defined as lacunae (blue) or vasculature (red) based on size and shape. B) Total porosity varies significantly with strain and sex, with a strain-sex interaction. B6 have a large sexual dimorphism, with B6 M having 4.1 times greater porosity than the least porous strain (CAST F). C) The median lacunar volume for each sample was determined based on all lacunae in the FOV. Lacunar volume varied significantly with strain and sex, with a strain-sex interaction. B6 M have 1.8 times larger lacunae than CAST F. D) The aspect ratio of each lacuna in the FOV was analyzed as a measure of elongation. An aspect ratio of 0 corresponds to a straight line while 1 is a perfect circle. Ellipses along the y-axis represent aspect ratios of 0.15 and 0.30. Lacunar aspect ratio varies with mouse strain. 129S1 F have a 1.3 times lower aspect ratio (i.e., more elongated) than AJ M. E) Phi (angle from z-axis) was calculated for each lacuna. The standard deviation (SD) of phi represents how uniformly aligned the lacunae are, with a smaller value being more aligned. The uniformity in alignment varies with mouse strain. NZO F have a 1.5 times larger SD of phi than B6 F. F) Pearson’s correlation matrix of all the lacunar traits measured in the Inbred Founder population. Black boxes group traits that cluster using hierarchical clustering. G) Contributions of each lacunar trait to the first four principal components of a PCA analysis. Traits that correlate highly contribute similarly to explain the variance in the population. *p<0.05, F vs M in same strain; # p<0.05, F strain vs F B6; ^ p<0.05, M strain vs M B6 p<0.05

Lacunae are ellipsoidal in shape, with the long-axis being parallel to the direction of loading^30^, therefore we quantified the lacunar elongation and orientation in the Inbred Founders. Median lacunar aspect ratio, a measure of elongation, also varied significantly with mouse strain (Fig 5E). In addition, we evaluated uniformity of the primary axis of the lacunae within each bone. For each lacunae, the angle of the lacunar long axis off the z-axis (phi) was calculated. We compared the SD of phi between mice, due to variation in bone placement in the XRM. Smaller SD of phi values represents greater uniformity of the primary axis of the population of osteocyte lacunae. The uniformity of alignment varied significantly with mouse strain (Fig 5F). The lacunar elongation (aspect ratio) and uniformity of alignment (SD of phi) are positively correlated with each other and lacunar number density (Fig 5G), indicating that in bones where lacunae are more densely packed, the lacuna tend to be more elongated and more aligned with each other. The lacunar traits that cluster together also contributed similarly to explain the population variance (Fig 5H). In summary, all osteocyte lacunar morphology traits are dependent upon mouse strain.

### Mineral composition minimally varies between Inbred Founder mice

Bone composition was analyzed using Raman spectroscopy on the transverse cross-section of the femur midsection (Supp Fig S1). Crystallinity, a measure affected by mineral length and organization, varied significantly between mouse strains (Fig 6A). The carbonate:phosphate ratio, which varies with architecture, age, and crystallinity^26, 31^, did not significantly vary between mouse strains (Fig 6B). One measurement of mineral:matrix (phosphate:proline) also varied significantly between mouse strains (Fig 6C). However, when mineral:matrix was measured using phosphate:amideIII, there was no significant variation (Fig 6D). In summary, the crystalline structure of the mineral varies modestly between Inbred Founders strains but bone composition is mainly conserved.

**Figure 6:**
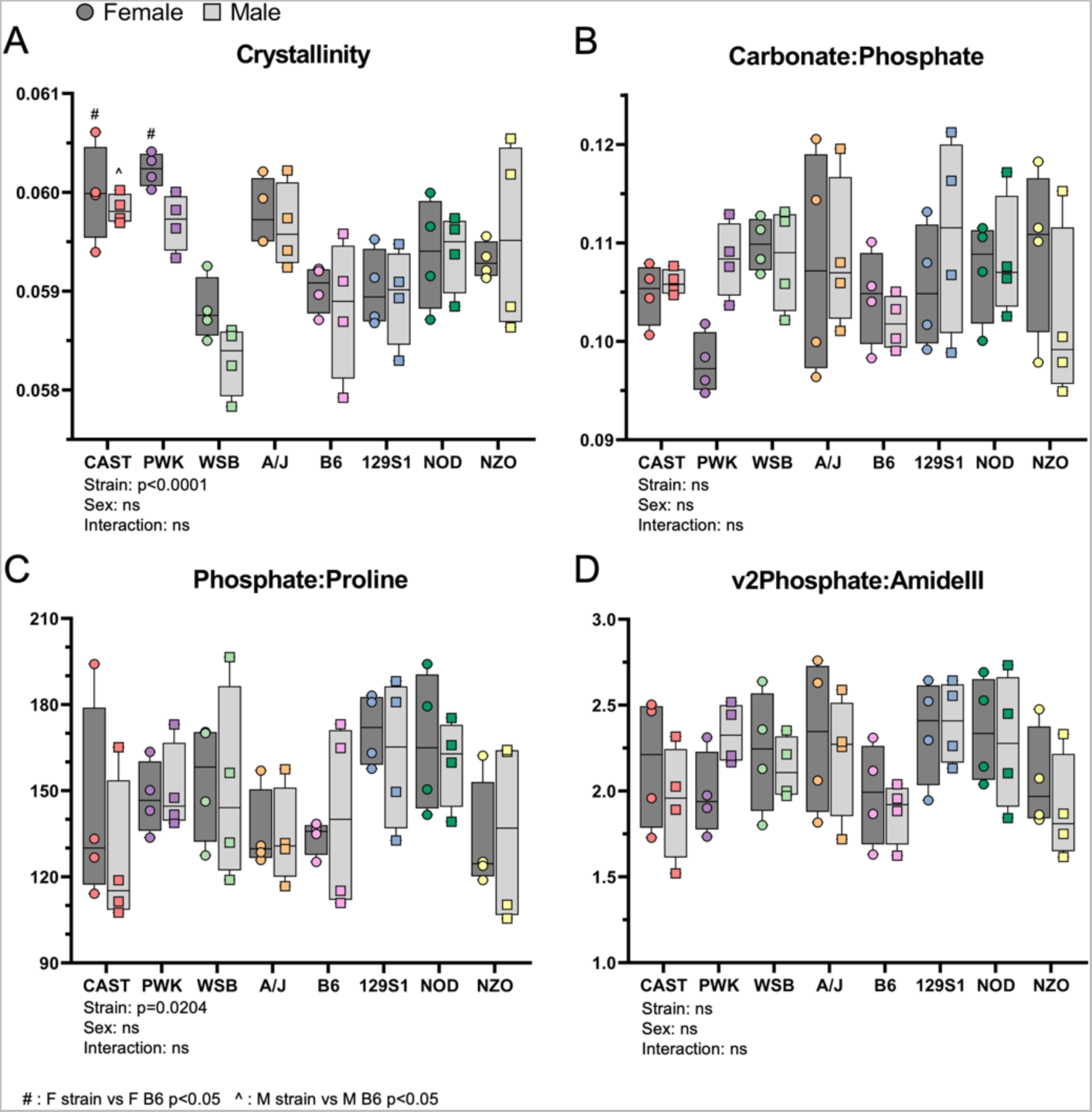
Raman spectroscopy was used to evaluate bone mineral composition. A) Crystallinity and B) Phosphate:Proline ratio vary significantly between strains. C) Carbonate:Phosphate ratio and D) vsPhosphate:amideIII ratio are not significantly different between strains.

### Multi-scale cortical bone traits are moderately to highly heritable in the Inbred Founder population

Broad-sense heritability was calculated for traits measured in the Inbred Founder strains using variance calculated from the two-factor ANOVA (Table 2). BMC had the highest heritability of 99.3% (H^2^ = 0.993) indicating almost all the variance in the population is attributed to genetic differences. In general, lacunar traits had lower heritability, but a few lacunar traits had very high levels of heritably, such as lacunar number density (H^2^ = 0.966). Heritability values were also similar between long bones. For the radius, femur, and tibia, cortical thickness and cortical area were the most heritable morphology traits. The rank order of µCT morphology traits in the femur and tibia are identical, with the femur always having a slightly higher value. Overall, of the 43 whole-body, whole-bone (radius, tibia, and femur), tissue, and lacunar traits, 35 (∼80%) had a heritability above 60% which is similar or greater than the reported heritability of BMD in humans^4–6^.

Whole-body mass significantly contributes to trait differences, but variation due to mouse strain remains significant Many of the bone traits we analyzed correlated with body weight (Fig 4A). There data were re-analyzed using an ANCOVA with body mass as a covariate to test if the effects of mouse strain remained after accounting for body mass. While body mass was a significant covariate for 22 out of 42 traits measured, the significant effect due to strain was maintained for 39/40 (98%) traits that were significant before body mass adjustment (Supp Table 1). PYD was the only trait that lost significance for strain after adjusting for body mass, but the significance of the strain-sex interaction was maintained. Notably, all lacunar level traits did not have weight as a significant covariate. To evaluate the effect of body weight on the separation of strains in the PCA, the PCA was re-run with body-weight adjusted traits. After adjustment, Inbred Founders still separated by strain with even more distinct clustering for PWK and CAST strains when compared to the PCA using unadjusted values (Supp Fig. S7B). After body mass adjustment, CAST radii more closely resemble radii of much larger mice (129S1) than from mice of similar size (PWK). NZO and B6 still fully separate from the other inbred strains. Additionally, how traits contribute to each principal component mainly remains the same after body-weight adjustment (Supp Fig S7 C,D). Heritability was also recalculated after body weight adjustment. The heritability of all traits except BMC, lacunar sphericity, and tibial cortical thickness increased or stayed the same after adjustment (Supp Table 2 and Supp Table 3). The lowest heritability value after body mass adjustment was 77.4% (phosphate:amideIII ratio). Therefore, while weight is an important factor when considering differences in skeletal traits between animals, the differences between mouse strains does not exclusively depend on mouse size (body mass).

## Discussion

Using two mouse models of genetic diversity we show that all measured cortical bone traits, from the whole-body to the osteocyte-lacunar length scale, vary with genetic background and are heritable. In the radius, whole-bone traits (morphology and mechanical properties) vary with both mouse strain (genetic background) and sex. Tissue level traits (tissue mineral density, material properties, and bone composition) vary with genetic background but not sex. Differences between inbred mouse strains for single traits are bone dependent (i.e., may be different between femur, radius, and tibia). Comparing the two populations, the Diversity Outbred (DO) mice are on average larger and protected from hyperglycemia. The DO population also has larger, stronger bones, but also has bone-dependent differences in tissue level properties. When all whole-body and radial traits are evaluated together, the DO population occupies a subset of the principal component space defined by the Inbred Founders although many DO mice do not resemble a single inbred strain, implying that complex gene interactions determine skeletal traits. Overall, genetic background significantly contributes to cortical bone phenotype, which indicates genetic control of bone traits across length scales.

We posed three hypotheses, which were tested using the two populations of genetically diverse animals. Our data support the first hypothesis that cortical bone traits vary with genetic background, consistent with a previous report that compared inbred mice^32^. Of the 43 traits we measured in the Inbred Founder strains, only two (carbonate:phosphate and phosphate:amideIII ratios) did not have strain as a significant factor. Additionally, broad-sense heritability, which represents the proportion of variability due to genetic difference, was greater than 20% for all traits, and after adjusting for body size heritability was greater than 77% for all traits. These values are higher than those reported by Al-Barghouthi et al^12^, who determined heritability values in DO mice as low as 12% after covariate adjustment. The mice used in that study were 12-weeks old, so there could be more non-genetic variation due to different rates of skeletal growth in mice that were not yet skeletally mature.

Furthermore, in the current study heritability was calculated using data from the eight Inbred Founder strains, where each group has genetically identical replicates reducing the intra-strain variation. The greatest variation we found between strains was for morphology and mechanical properties (e.g. cortical area and ultimate force). These traits tend to have higher heritability values, larger fold-differences between high and low groups, and contribute to the first principal component (PC1) of the PCA, which explains 55% of population variation. This same trend was reported in Al-Barghouthi et al., where morphology traits had the highest heritability values^12^. In contrast, we observed that the tissue-level properties (e.g. ultimate stress, TMD, and mineral:matrix) tend to have less variability between strains and more variability within a strain. These traits have lower fold-differences and mainly contribute to PC2, which explains only about 20% of the population variance. Overall, these results indicate that bone material properties are fairly conserved between strains, whereas how this bone is distributed varies markedly between strains.

To our knowledge, this is the first study to quantify osteocyte lacunar morphology across different mouse strains, as all previous work has been done on mice from a C57BL/6 background. At the cellular length scale, lacunar size traits (e.g., diameter, volume) tend to have moderate heritability (H^2^ = 0.38-0.63). By contrast, the number of lacunae (e.g., lacunar number density) and their shape (e.g., sphericity, aspect ratio) have relatively high heritability (0.76-0.97). Therefore, the shape of the lacunae and how densely they are packed are strongly determined genetically in the Inbred Founder mice, whereas the volume of a single lacunae is less so.

Contrary to our second hypothesis, many traits of the DO population have different means and smaller ranges than the Inbred Founder population. For 24 out of the 29 traits measured in both Inbred Founders and DO mice, there was a significant difference in mean values between the populations. As a population, the DO had larger skeletal size (BMC), body weight, and healthier glucose levels than Inbred Founders. For the radius, tibia, and femur, the DO bones were larger (e.g., greater cortical area), although medullary area was not different. The DO had higher mechanical properties, but diminished material properties. The combination of higher total area but lower tissue mineral density seen in the DO population matches the preferred bone trait set established in Jepsen et al.^33^, and resulted in greater ultimate force. The range of values for each trait in DO mice was generally lower than in Founders, and extreme values were eliminated. For example, for each µCT parameter, the DO have a smaller coefficient of variation (CV) compared to the Inbred Founders. For whole-body traits, both maximum and minimum extreme values were eliminated in the DO. For morphology traits, only minimum extremes were eliminated; there were no DO mice with bones as small as the smallest Inbred Founder bones, however there were DO mice with bones as large as the largest Inbred Founder bones. When the DO mice were plotted onto the PCA space defined by the Inbred Founders, the DO population resided in a subset, hybrid space emphasizing the elimination of extreme values and the mixing of phenotypes. The elimination of extreme values in the outbred population implies epistasis in the Inbred Founder strain with the extreme phenotypes, since disruption of the exact allele combination removes the phenotype.

While other groups have reported bone phenotypes for either the Inbred Founders^15^ or the DO mice^12^ no one has directly compared the two populations. Turner et al.^34^ compared two inbred strains (B6 and C3H) to multiple recombinant inbred lines (BXH RI) and showed that none of the measured traits grouped the RI strains into subsets resembling either of the two progenitors. Additionally, none of the RXH RI strains had ultimate force values approaching the high-strength C3H femurs, supporting epistasis of bone traits as our data suggest. In contrast, Jepsen et al.^33^ compared A/J and B6 inbred strains to multiple recombinant inbred lines (AXB/BXA RI) and showed many RI lines with femurs smaller than A/J and larger than B6. Depending on the alleles available and the specific combination per animal, bone phenotypes vary considerably, indicating that complicated interactions between genomic regions control skeletal phenotypes.

In support of our third hypothesis, many relationships between traits seen in the Inbred Founder population are maintained in the DO population, especially those between morphology and mechanical properties. However, correlations between most traits and body weight or bone length were greatly diminished. In the Inbred Founder strains, a relatively large mouse (by body weight) would typically have a large bone. However, in the DO mice, this relationship is disrupted. For example, the correlation between cortical area and weight is non-significant in the DO population (r = 0.75 for the Inbred Founders, r = 0.23 in the DO population). On the other hand, across both populations, one of the strongest correlations is between bone area and ultimate force, with larger bones being stronger. In fact, we observed an almost identical relationship (slope and intercept are not significantly different) between cortical area and ultimate force in the two populations of genetic diversity, suggesting that this relationship is conserved in all *Mus musculus*. Previously, we evaluated the relationships between traits in a large population of advanced intercross mice (F34 LGXSM AI), which have a large range of body weight and bone size^29^. We see many similar relationships between traits in that study as we discovered herein. Specifically, in both studies morphology traits are highly correlated with each other expect for cortical thickness, which only has low correlations with other traits. Morphology traits also highly correlate with stiffness and strength but not brittleness (PYD). All material properties are highly correlated with each other but relatively independent of morphology. The current study provides additional support that our proposed reduced set of parameters (Ct.Ar, Ma.Ar, Ult.Force, PYD, Ult. Stress)^29^ are useful to describe variation in mouse long bones from any population. Jepsen et al. compared femurs from multiple RI lines, and showed a complex, biologically important relationship between bone morphology and tissue quality^33^. Specifically, there was a positive correlation between tissue mineral density and cortical thickness in the femurs in that study, which holds true for the radii in both the Inbred Founder population and the DO population herein. Thus, our data add support to the idea that there is a coordinated, biological regulation of the relative rates of periosteal and endocortical expansion (which determine cortical thickness) and tissue mineral density^33^.

Many measured traits correlated significantly with body weight, which had a 4.5-fold range from smallest to largest strain of the Inbred Founders . Therefore, we asked whether differences in traits between strains were due solely to differences in body size. After adding body weight as a covariate to the ANOVA analysis, all traits maintained a significant strain or strain/sex effect. The clustering of inbred strains on the PCA also became more distinct and separate after body weight adjustment. Additionally, the heritability increased for most traits after body weight adjustment. This analysis indicates that variations in body size alone do not explain variations in bone traits, and indicates that there are genes that independently control bone traits and body size.

We acknowledge some limitations of our study. First, while we were able to collect data from 188 individual animals, some strain/sex groups had small sample size. Although 9 mice per strain per sex were ordered, we ended up with only 4 female CAST mice due to attrition of this difficult-to-handle strain. Additionally, only 4 samples per sex per strain were evaluated for lacunar (XRM) and bone composition (Raman spectroscopy) analysis. The interpretation from these outcomes, especially those with high variability within a strain/sex group should be supported by more data. When comparing the Inbred Founder mice to the Diversity Outbred population, only 25 DO animals per sex were sampled. We are confident this sample size provides a reasonable estimate of the population mean and distribution, but it is possible adding more DO animals would alter the differences we observed. Second, because two of the inbred strains (NOD, NZO) are commonly used as models for diabetes, we tracked the glycemic status of our mice. Only two groups consistently became hyperglycemic (NOD female and NZO males). No mice were treated with insulin or other means to manage their hyperglycemia, in order to avoid any influence of treatments on bone traits. For NOD and NZO mice, most traits did not exhibit a large sex difference even though hyperglycemia only occurred in one sex by 22 weeks old, which suggests that these traits were not significantly altered by high glucose levels in our study.

In summary, we find that multi-scale cortical bone traits vary with genetic background and are moderately to highly heritable. While we noted significant variations in bone composition, these were less pronounced than variations in bone morphology, indicating that the bone composition between mice is well conserved while bone distribution varies widely. There are several novel findings from this study. First, this is the first study to report osteocyte lacunar traits for genetically diverse mice and to show that these traits are genetically regulated. Second, this is the first study to directly compare bone phenotypes between the DO mice and their inbred progenitors (Inbred Founders). This provides insight into how traits change with outbreeding. Lastly, this is the first study to evaluate the conservation of intra-bone relationships in multiple genetically diverse populations, providing support for how robust these relationships are for *Mus musculus*. This work supports future use of these genetically diverse populations to discover novel genes contributing to cortical bone traits, especially at the lacunar length scale.

## Supporting information

All Supplemental Information

All Data for Diversity Outbred Mice

All Data for Inbred Founder Mice

## Acknowledgments

We thank members of the Flores Lab (Dr. Katharine Flores, Dr. Porter Weeks, and Devin Williamson) for allowing us to use the automatic polisher for our MMA blocks. We thank Dr. Jeffry Nyman and Dr. Rafay Ahmed for advice and input with our Raman Spectroscopy analysis code. We thank the Materials and Instrumentation and Multimodal Imaging Core Facility (MIMIC) at the University of Colorado (RRID: SCR 019307) for completing the Raman Spectroscopy. We thank the Washington University Center for Cellular Imaging (WUCCI) for use of the XRM (S10 OD021694) and the Musculoskeletal Research Center (MRC -NIH P30 AR074992) for support of microCT and mechanical testing resources. Finally, we thank our funding sources (NIH / NIAMS R01 AR047867, NIH T32 DK108742, NIH T32 EB018266, NIH R21 079052, NIH R01 AR071657, Jax DO Grant).

