## Supplementary material for "Multi-Scale Cortical Bone Traits Vary in Two Mouse Models of Genetic Diversity": All Supplemental Information

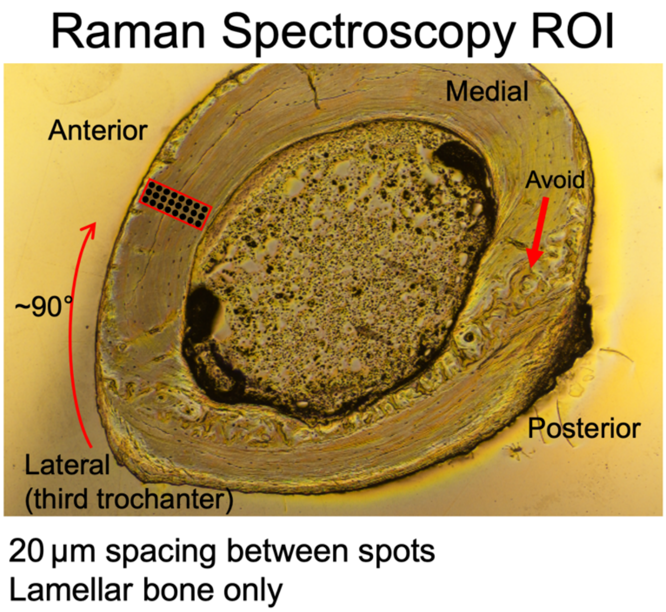


*Supp Figure S1: Region of interest for Raman spectroscopy on the transverse cross-section of the femur. Bones were embedded in PMMA and polished to a smoothness of 0.05 µm. The red box shows the region where measurements were taken.*

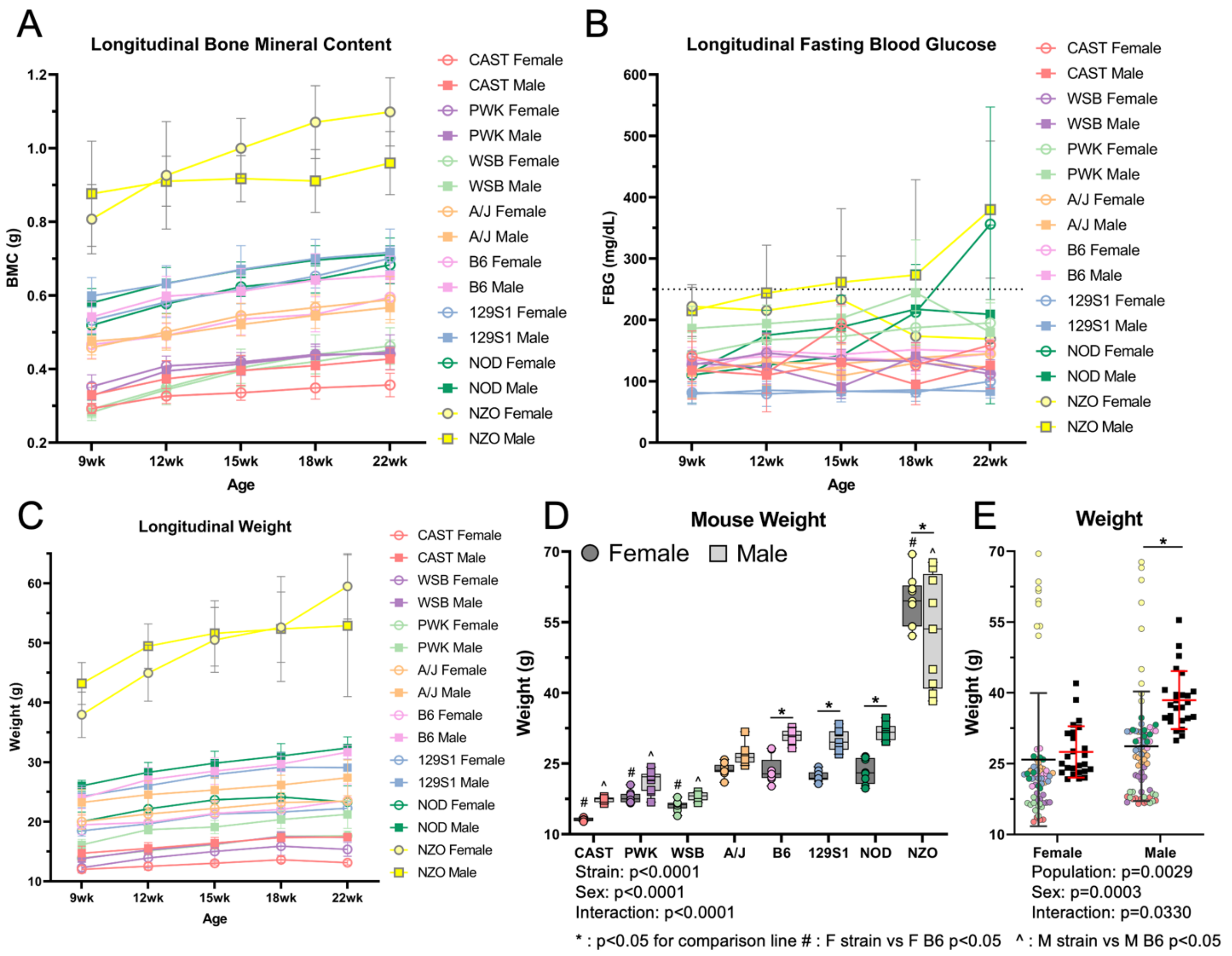


*Supp Figure S2: A) BMC of founder strain mice overtime showing healthy skeletal growth. BMC has plateaued around 22wks of age, with no significant differences between 18wk and 22wk values for any group. B) FBG of founder strains overtime showing the onset of hyperglycemia. NOD F become hyperglycemic between 18wk and 22wk of age. NZO M become hyperglycemic earlier, between 12wk and 15wk of age. C) Trends in body weight match those of BMC. NOD F begin to lose weight around the onset of hyperglycemia. D) At 22wks of age, the Inbred Founder have a significant variations due to strain, sex, and a sex-strain interaction. E) At 22wks the DO mice have a higher weight with less variation than the Inbred Founder population. * : p<0.05 for comparison line # : F strain vs F B6 p<0.05 ^ : M strain vs M B6 p<0.05*

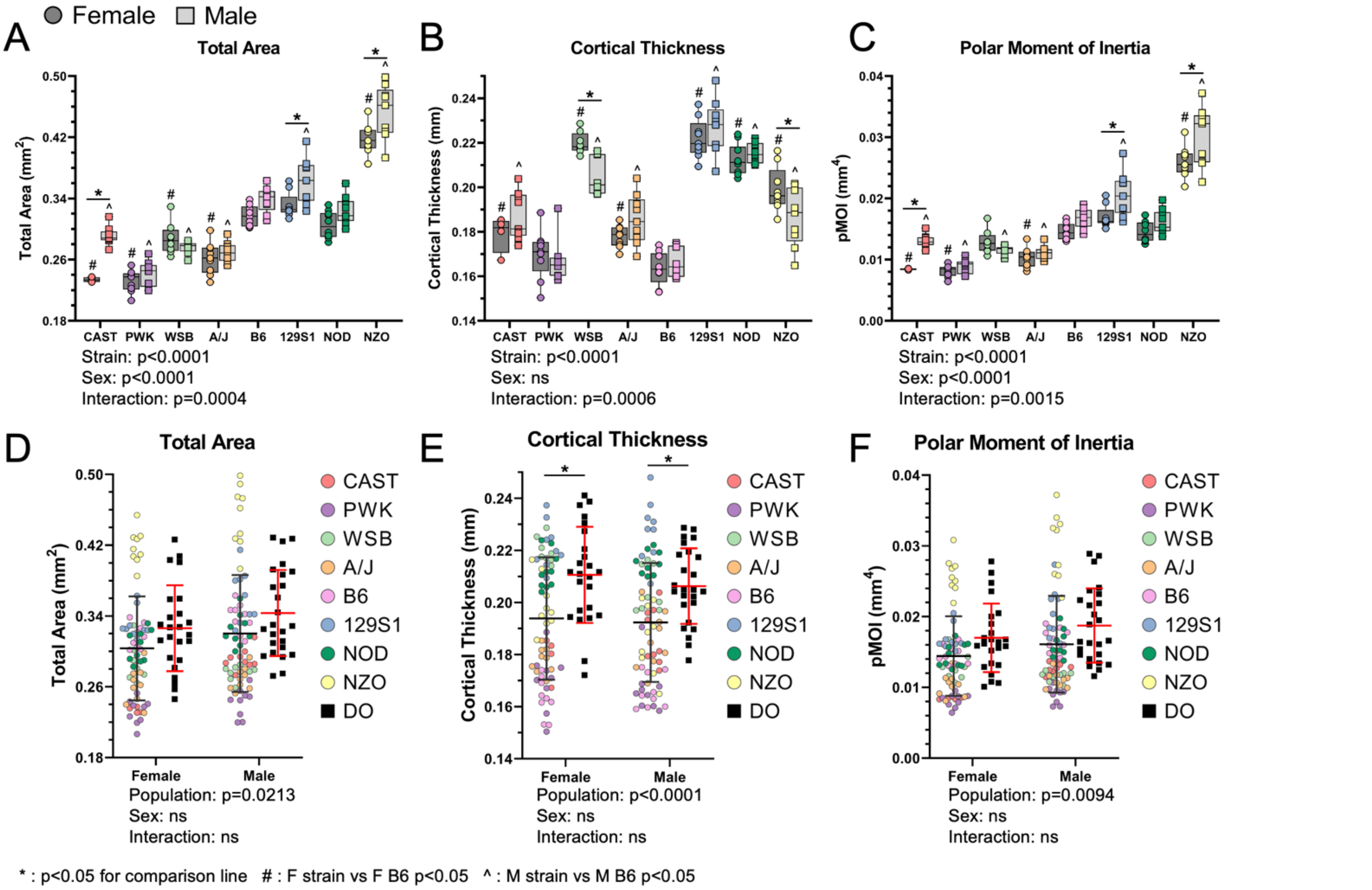


*Supp Figure S3: Radial morphology changes with strain in a sex dependent manner. Additional radial traits not reported in Fig 2. * : p<0.05 for comparison line # : F strain vs F B6 p<0.05 ^ : M strain vs M B6 p<0.05*

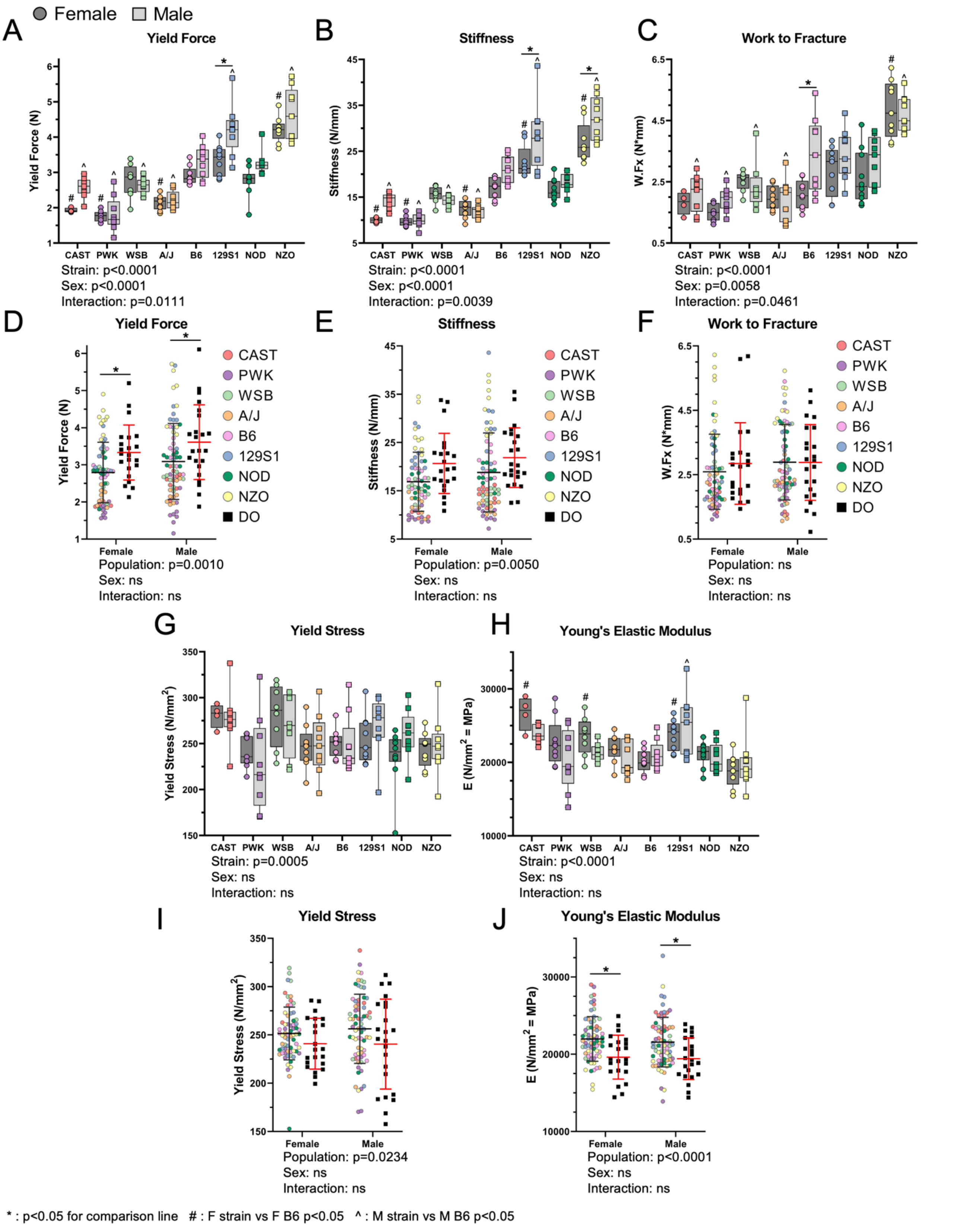


*Supp Figure S4: Radial material and mechanical properties changes with strain in a sex dependent manner. Mechanical (A-F) and material properties (G-J) not reported in Fig 3. * : p<0.05 for comparison line # : F strain vs F B6 p<0.05 ^ : M strain vs M B6 p<0.05*

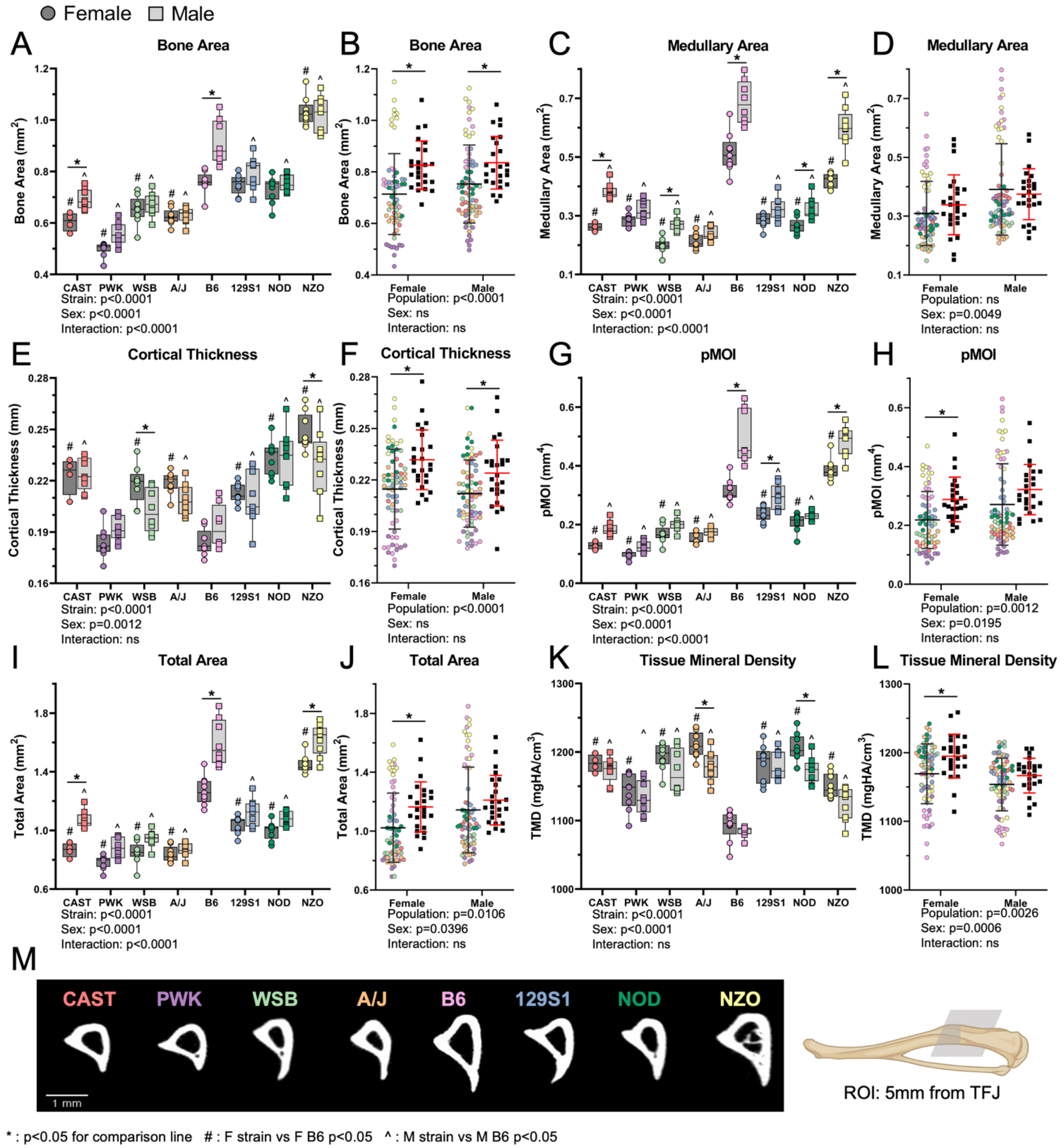


*Supp Figure S5: Tibial morphology changes with strain in a sex dependent manner. Representative cross-sections 5mm proximal of the tibia-fibula junction (TFJ) show the variation in bone shape and size between the eight inbred founder strains. * : p<0.05 for comparison line # : F strain vs F B6 p<0.05 ^ : M strain vs M B6 p<0.05*

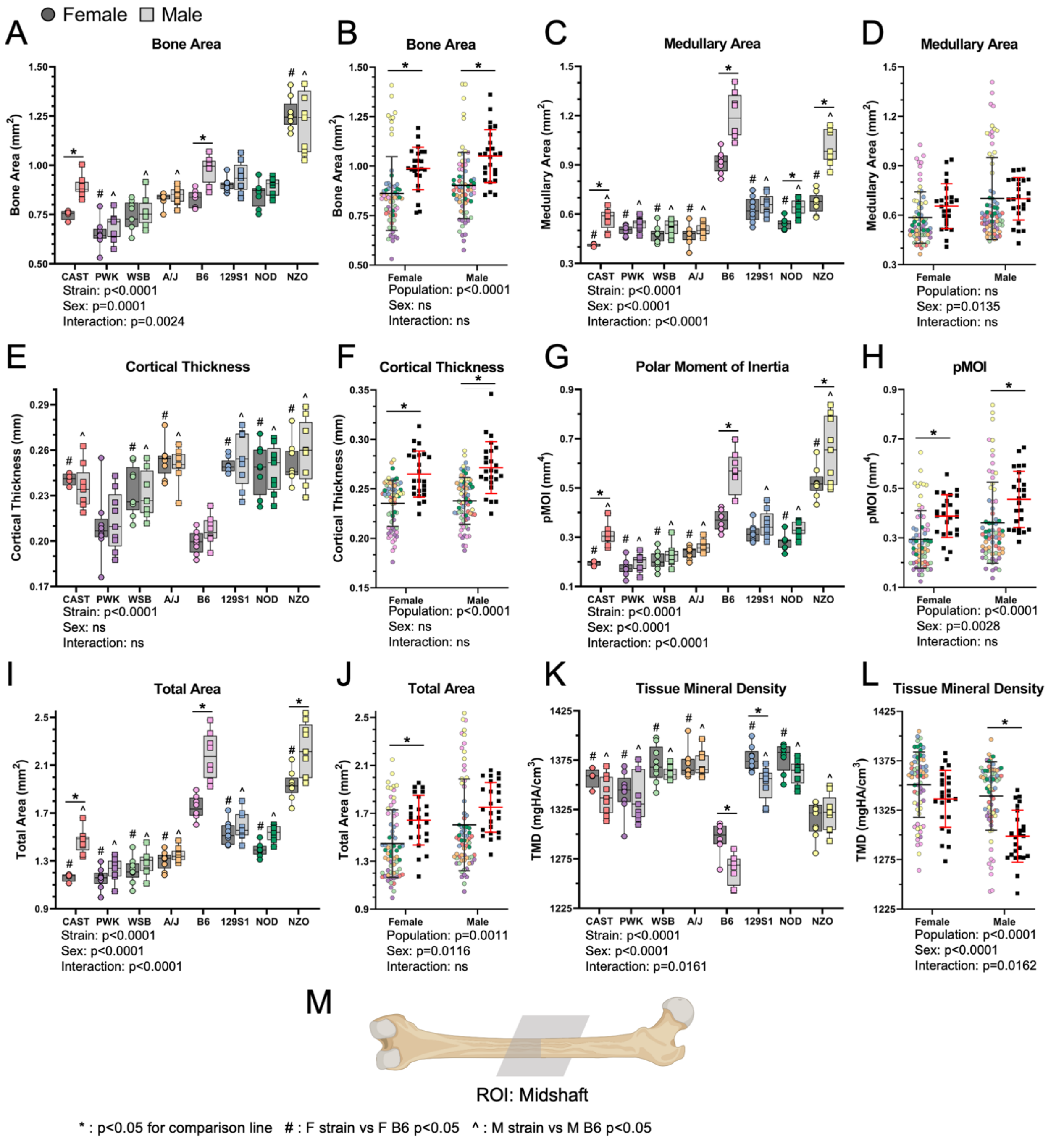


*Supp Figure S6: Femoral morphology changes with strain in a sex dependent manner. * : p<0.05 for comparison line # : F strain vs F B6 p<0.05 ^ : M strain vs M B6 p<0.05*

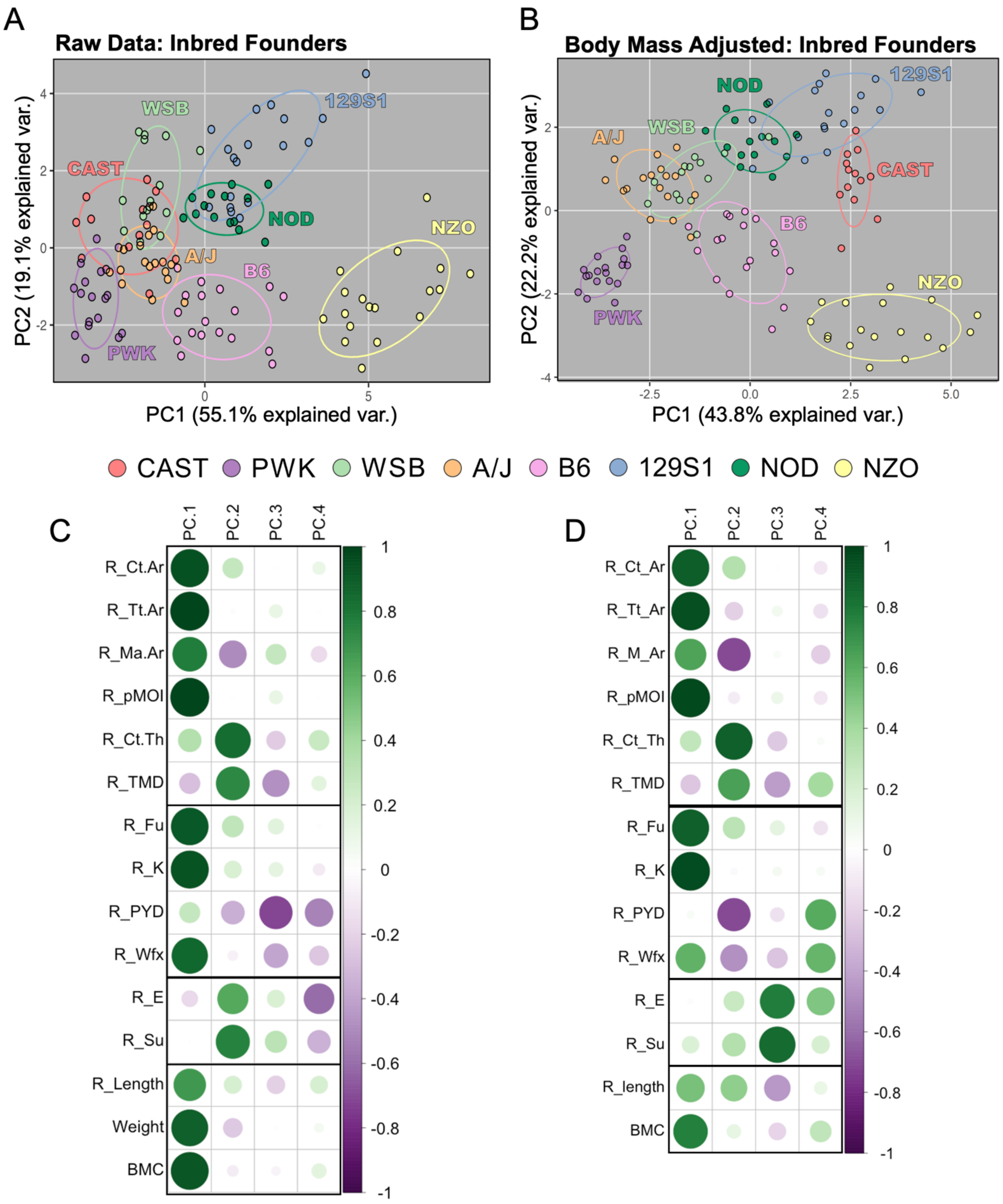


*Supp Figure S7: PCA of raw and body mass adjusted data from Inbred Founder mice. A) PCA of raw data repeated from Fig4C for ease of contrast. B) the Inbred Founder strains cluster more tightly after body weight adjustment. Contributions of each trait to the first four principal components for the C) raw data and D) body mass adjusted data.*

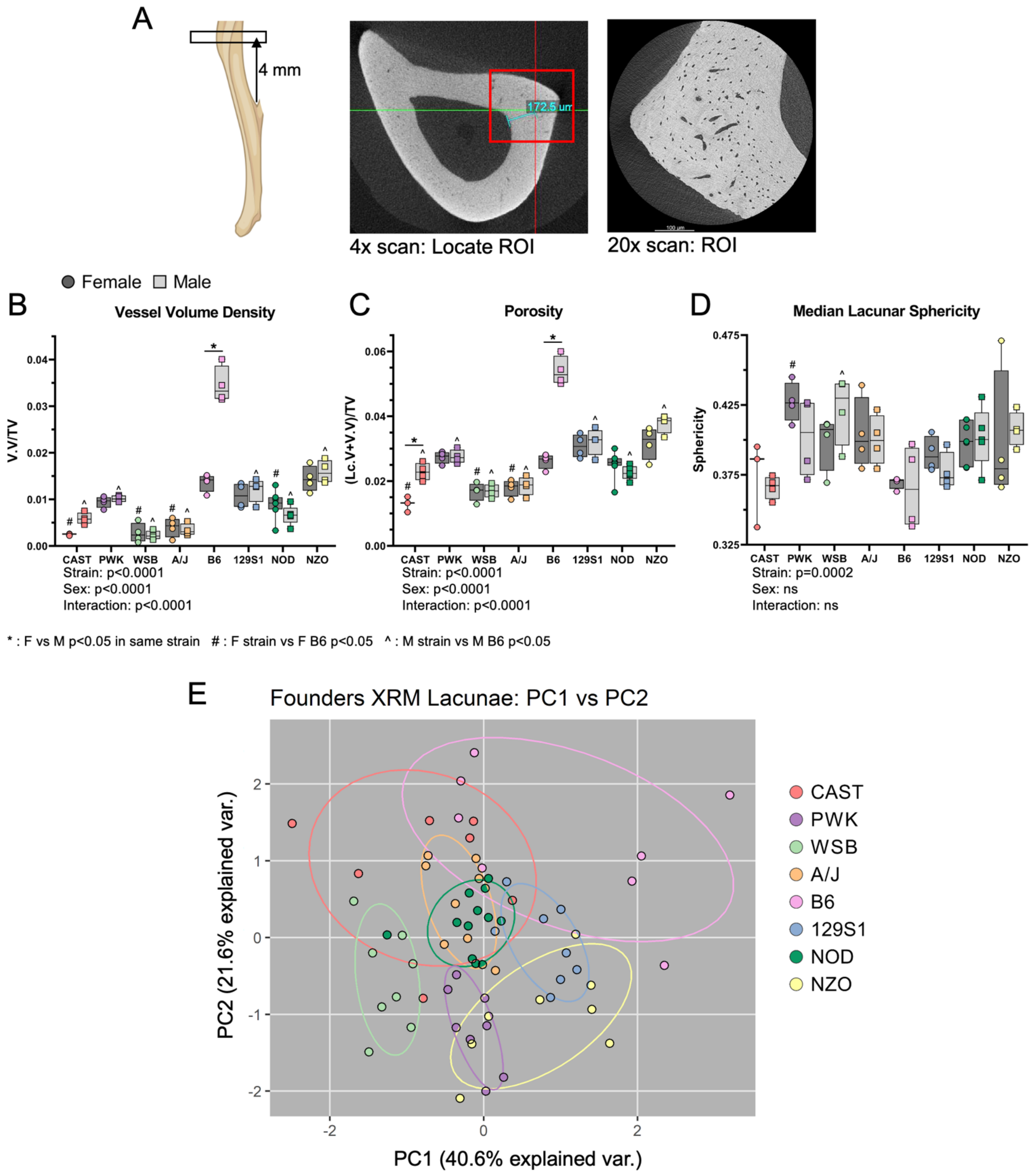


*Supp Figure S8: A) Schematic showing the region of interest scanned and analyzed. The scanning region of approximately 500 µm x 500 µm x 500 µm centered 4 mm proximal to the distal TFJ at the postero-lateral apex. B-D) XRM outcomes not reported in Fig 5. E) PCA using all the lacunar traits shows the Inbred Founder strains do not clearly separate. * : F vs M p<0.05 in same strain # : F strain vs F B6 p<0.05 ^ : M strain vs M B6 p<0.05*

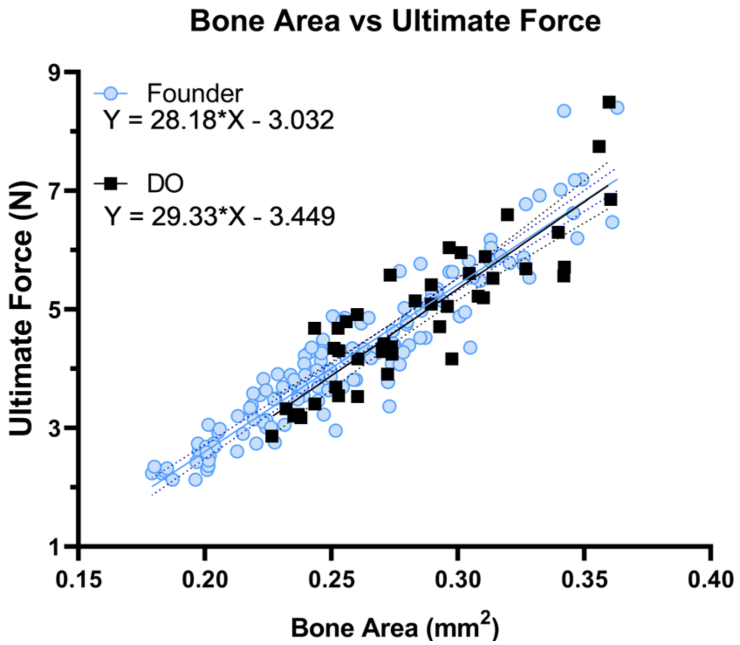


*Supp Figure S9: Bivariate plot highlighting how the relationship between cortical area and ultimate force is highly conserved. The slopes and intercepts between the two linear-regression lines are not significantly different.*

*Supp Table 1: ANCOVA p-value results with body weight as a co-factor.*

| | | Factor p-value | | | |
|---|---|---|---|---|---|
| | Trait | Weight | Strain | Sex | Strain*Sex |
| Radius | Cort. Area | **0.001** | **0.001** | 0.181 | **0.001** |
| | Total Area | **0.001** | **0.001** | **0.001** | **0.001** |
| | Med. Area | 0.609 | **0.001** | **0.001** | **0.001** |
| | Cort. Thickness | **0.001** | **0.001** | **0.029** | **0.022** |
| | pMOI | **0.001** | **0.001** | **0.005** | **0.001** |
| | TMD | 0.381 | **0.001** | 0.881 | 0.398 |
| Radius | Stiffness (K) | **0.001** | **0.001** | **0.044** | **0.001** |
| | Ult. Force | **0.001** | **0.001** | 0.227 | **0.001** |
| | Yield Force | **0.001** | **0.001** | **0.011** | **0.004** |
| | PYD | 0.414 | 0.266 | **0.010** | **0.022** |
| | Wfx | 0.128 | **0.001** | 0.070 | 0.219 |
| | Ult. Stress | **0.005** | **0.001** | **0.013** | **0.043** |
| | Yield Stress | **0.011** | **0.001** | 0.450 | 0.520 |
| | Elastic Modulus | **0.016** | **0.001** | **0.008** | 0.065 |
| Tibia | Lc.Vol Density | 0.233 | **0.001** | **0.001** | **0.001** |
| | Vess.Vol Density | 0.805 | **0.001** | **0.001** | **0.001** |
| | Porosity | 0.619 | **0.001** | **0.001** | **0.001** |
| | Lc.No Density | 0.837 | **0.001** | 0.942 | **0.032** |
| | Lc.Vol | 0.189 | **0.001** | **0.001** | **0.001** |
| | Lc.AspectRatio | 0.432 | **0.001** | 0.074 | 0.507 |
| | Lc.SD.Phi | 0.196 | **0.004** | 0.289 | 0.529 |
| | Lc.Vol/SA | 0.252 | **0.002** | **0.031** | **0.040** |
| | Lc.Diameter | 0.300 | **0.001** | **0.008** | **0.034** |
| | Lc.Sphericity | 0.659 | **0.001** | 0.590 | 0.579 |
| Femur | Phos:Proline | 0.915 | **0.028** | 0.634 | 0.985 |
| | v2Phos:AmideIII | 0.568 | 0.086 | 0.726 | 0.670 |
| | Carb:Phos | 0.361 | 0.325 | 0.823 | 0.564 |
| | Crystallinity | 0.378 | **0.001** | 0.093 | 0.642 |
| Body | BMC | **0.001** | **0.001** | **0.001** | **0.025** |
| | FBG | **0.001** | **0.001** | 0.070 | **0.001** |
| Tibia | Cort. Area | **0.001** | **0.001** | **0.003** | **0.001** |
| | Total Area | **0.001** | **0.001** | **0.001** | **0.001** |
| | Med. Area | 0.471 | **0.001** | **0.001** | **0.001** |
| | Cort. Thickness | **0.001** | **0.001** | **0.001** | **0.031** |
| | pMOI | **0.001** | **0.001** | **0.001** | **0.001** |
| | TMD | 0.179 | **0.001** | **0.001** | 0.367 |
| Femur | Cort. Area | **0.001** | **0.001** | 0.399 | **0.005** |
| | Total Area | **0.001** | **0.001** | **0.001** | **0.001** |
| | Med. Area | **0.017** | **0.001** | **0.001** | **0.001** |
| | Cort.Thickness | **0.036** | **0.001** | 0.738 | 0.596 |
| | pMOI | **0.001** | **0.001** | **0.001** | **0.001** |
| | TMD | 0.517 | **0.001** | **0.001** | 0.098 |

*Supp Table 2: Heritability of all traits reported in main paper (left is repeated from Table 2) (right is body-weight adjusted)*

**Raw Data: Inbred Founders**

| Trait | Heritability ($H^2$) |
|---|---|
| BMC | 0.993 |
| Cortical Thickness | 0.985 |
| TMD | 0.967 |
| Lc.No Density | 0.966 |
| FBG | 0.910 |
| Lc.Sphericity | 0.887 |
| Cortical Area | 0.884 |
| Crystallinity | 0.844 |
| Weight | 0.840 |
| Ult. Force | 0.836 |
| Lc.SD.Phi | 0.835 |
| V.Vol Density | 0.827 |
| Phos:Proline | 0.806 |
| pMOI | 0.799 |
| Total Area | 0.799 |
| Stiffness (K) | 0.776 |
| Lc.AspectRatio | 0.755 |
| Ult. Stress | 0.746 |
| Work-to-Fx | 0.741 |
| Phos:AmideIII | 0.741 |
| Medullary Area | 0.722 |
| Yield Force | 0.718 |
| Yield Stress | 0.717 |
| Porosity | 0.689 |
| Modulus (E) | 0.684 |
| Lc.Diameter | 0.634 |
| Carb:Phos | 0.574 |
| Lc.Vol/SA | 0.535 |
| Lc.Vol Density | 0.516 |
| Lc.Vol | 0.383 |
| PYD | 0.209 |

**Body Mass Adjusted: Inbred Founders**

| Trait | Heritability ($H^2$) |
|---|---|
| BMC | 0.800 |
| Cortical Thickness | 0.993 |
| TMD | 0.973 |
| Lc.No Density | 0.975 |
| FBG | 0.936 |
| Lc.Sphericity | 0.881 |
| Cortical Area | 0.992 |
| Crystallinity | 0.873 |
| Weight | NA |
| Ult. Force | 0.984 |
| Lc.SD.Phi | 0.835 |
| V.Vol Density | 0.969 |
| Phos:Proline | 0.806 |
| pMOI | 0.960 |
| Total Area | 0.989 |
| Stiffness (K) | 0.983 |
| Lc.AspectRatio | 0.871 |
| Ult. Stress | 0.983 |
| Work-to-Fx | 0.958 |
| Phos:AmideIII | 0.774 |
| Medullary Area | 0.959 |
| Yield Force | 0.968 |
| Yield Stress | 0.897 |
| Porosity | 0.975 |
| Modulus (E) | 0.962 |
| Lc.Diameter | 0.775 |
| Carb:Phos | 0.866 |
| Lc.Vol/SA | 0.780 |
| Lc.Vol Density | 0.969 |
| Lc.Vol | 0.893 |
| PYD | 0.804 |

| |
|---|
| Whole Body |
| Whole Bone |
| Tissue Level |
| Cellular Level |

*Supp Table 3: Heritability of uCT parameters from all three bones. Radial heritability values are repeated from Table 2. Right side shows heritability for body weight adjusted values.*

**Raw Data: Inbred Founders**

| Trait | Bone | Heritability ($H^2$) |
|---|---|---|
| Ct.Th | Radius | 0.985 |
| TMD | Radius | 0.967 |
| Ct.Th | Femur | 0.937 |
| Ct.Th | Tibia | 0.886 |
| Ct.Ar | Radius | 0.884 |
| Ct.Ar | Femur | 0.861 |
| Ct.Ar | Tibia | 0.807 |
| pMOI | Radius | 0.799 |
| Tt.Ar | Radius | 0.799 |
| TMD | Femur | 0.741 |
| Ma.Ar | Radius | 0.722 |
| TMD | Tibia | 0.636 |
| Tt.Ar | Femur | 0.621 |
| Tt.Ar | Tibia | 0.614 |
| pMOI | Femur | 0.607 |
| pMOI | Tibia | 0.597 |
| Ma.Ar | Femur | 0.522 |
| Ma.Ar | Tibia | 0.503 |

**Body Mass Adjusted: Inbred Founders**

| Trait | Bone | Heritability ($H^2$) |
|---|---|---|
| Ct.Th | Radius | 0.993 |
| TMD | Radius | 0.973 |
| Ct.Th | Femur | 0.956 |
| Ct.Th | Tibia | 0.870 |
| Ct.Ar | Radius | 0.992 |
| Ct.Ar | Femur | 0.991 |
| Ct.Ar | Tibia | 0.991 |
| pMOI | Radius | 0.960 |
| Tt.Ar | Radius | 0.989 |
| TMD | Femur | 0.969 |
| Ma.Ar | Radius | 0.959 |
| TMD | Tibia | 0.917 |
| Tt.Ar | Femur | 0.957 |
| Tt.Ar | Tibia | 0.968 |
| pMOI | Femur | 0.909 |
| pMOI | Tibia | 0.947 |
| Ma.Ar | Femur | 0.837 |
| Ma.Ar | Tibia | 0.926 |

| |
|---|
| Radius |
| Femur |
| Tibia |
